# FAK inhibitor treatment systemically regulates the glioblastoma immune environment via blocking monocyte trafficking

**DOI:** 10.1101/2025.11.26.690731

**Authors:** Emily R Webb, Annabel Black, Giovana Carrasco, Muhammad Furqan, Robert L Hollis, Alexander E P Loftus, Julián Corzo Ochoa, Romain Enjalbert, Tara Best, Bo Peng, Morwenna Muir, Fraser Laing, Martin Lee, Sofian Al Shboul, Tongjie Wang, Colin Smith, Ted R Hupp, Ajitha Rajan, Javier A Alfaro, Paul M Brennan, Zhaoyuan Liu, Florent Ginhoux, Miguel O Bernabeu, Alan Serrels, Rebecca Gentek, Margaret C Frame, Valerie G Brunton

**Affiliations:** Cancer Research UK Scotland Centre, Institute of Genetics and Cancer, University of Edinburgh, Crewe Road South, Edinburgh, EH4 2RX, United Kingdom; Centre for Medical Informatics, Usher Institute, University of Edinburgh, Edinburgh, EH16 4UX, United Kingdom; Department of Pharmacology and Public Health, Faculty of Medicine, The Hashemite University, Zarqa, 13133, Jordan; School of Informatics, The University of Edinburgh, Edinburgh, United Kingdom; Centre for Clinical Brain Sciences, Institute for Neuroscience and Cardiovascular Research, University of Edinburgh, Edinburgh, EH8 9JZ, United Kingdom; Riddell Centre for Cancer Immunotherapy, Arnie Charbonneau Cancer Institute, Cumming School of Medicine, University of Calgary, Calgary, Alberta, Canada; Department of Biochemistry & Molecular Biology, Cumming School of Medicine, University of Calgary, Calgary, Alberta, Canada; Shanghai Institute of Immunology, Department of Immunology and Microbiology, Shanghai Jiao Tong University School of Medicine, Shanghai 200025, China; Innovations in Food & Chemical Safety Programme (IFCS), A*STAR; Singapore Immunology Network, 8A Biomedical Grove, Singapore 138648, Singapore; School of Biological Sciences, Nanyang Technological University, Singapore 637551, Singapore; Shanghai Institute of Immunology, Shanghai JiaoTong University School of Medicine, Shanghai, China; Translational Immunology Institute, SingHealth Duke-NUS Academic Medical Centre, Singapore, Singapore; The Bayes Centre, University of Edinburgh, Edinburgh EH8 9BT, United Kingdom; Institute for Regeneration and Repair, Centre for Reproductive Health, University of Edinburgh, EH16 4UU, United Kingdom

**Keywords:** Glioblastoma, FAK, monocytes, T cell exhaustion, monocyte derived macrophages, integrin adhesion, a4b1 integrin, CX3CR1, bone marrow

## Abstract

Glioblastoma (GBM) is the most common primary malignant brain tumour in adults with dismal survival rates, and current therapies, including most immunotherapies, are not efficacious due to the highly immunosuppressive microenvironment. Studies in other solid cancers report that impairment of the integrin effector pathway involving focal adhesion kinase (FAK) can promote anti-tumour immune responses. Therefore, we set out to address whether, and if so how, suppressing FAK function may influence GBM by using both tumour cell-specific FAK gene deletion and systemic delivery of a clinically relevant FAK kinase inhibitor (FAKi) VS-4718 in an orthotopic murine stem cell model of GBM. We found that treatment with the FAKi, but not tumour cell-specific FAK gene deletion, resulted in GBM clearance and improved survival. This was dependent on adaptive immunity, and tumour-infiltrating T cells in FAKi-treated tumours displayed increased cytotoxic potential and reduced exhaustion. We also found a significant reduction in immuno-suppressive peripherally-derived macrophages and FAKi treatment caused sequestration of inflammatory monocytes within the bone marrow, resulting in impaired monocyte trafficking to tumours as judged by adoptive transfer. This is due to suppression of key adhesion and migration signalling through α4β1 integrin and CX3CR1 in peripheral monocytes. Our work here describes a previously unidentified role for FAK in trafficking of peripheral suppressive macrophages to GBM tumours, reducing T cell exhaustion and promoting anti-tumour immunity. This highlights a new way in which systemic FAK inhibitors can be used to provide a beneficial immune modulatory strategy for the treatment of GBM.

## Introduction

Glioblastoma (GBM) is the most common subtype of diffuse gliomas, accounting for >50% of adult gliomas and 16% of all primary brain tumours, alongside being the most aggressive subtype with a median survival of 15 months with surgical resection ^1,2^. Despite an intense standard of care treatment regimen (surgical resection, radiotherapy and temozolomide chemotherapy), median survival rates remain poor, with no new effective treatments approved in decades ^3^. Immunotherapy has shown great promise in treating extra-cranial solid tumours; however, its efficacy in cranial tumours, particularly GBM, is poor ^4-6^. This apparent lack of efficacy in GBM is due to a number of factors resulting in an immunotherapy-averse environment including: intra-tumoural heterogeneity ^7,8^, shielding by the blood brain barrier (BBB) ^9^, and a tendency towards developing a strongly immunosuppressive tumour microenvironment ^10-12^. Despite this, some novel approaches to immunotherapy regimes have shown encouraging outcomes in certain GBM patients ^13-15^, particularly chimeric antigen receptor T cell therapy ^16^, indicating that it is possible to render GBM immunotherapy-responsive.

The key to tumour clearance in many cancers relies on a functioning anti-tumour immune response, enabling robust and sustained tumour cell killing by cytotoxic immune cells within the tumour microenvironment (TME) ^17,18^. However, within brain tumours, the TME is largely characterised by an accumulation of peripherally derived myeloid cells, accounting for around 50% of the immune cell mass within GBM ^10,11^, and their accumulation correlates with the aggressive mesenchymal subtype ^7^. Importantly, the majority of the infiltrating myeloid cells are immunosuppressive in nature ^19,20^. Together with a distinct lack of cytotoxic T cell infiltration ^10,21^, this results in an immunosuppressive TME within GBM, and goes some way to explaining the failure of many T cell directed immunotherapeutics in this disease setting ^2,5,6,22^. Therefore, to unlock robust anti-tumour immunity in GBM, novel ways of modulating the suppressive myeloid rich microenvironment are needed.

Over recent years, the integrin adhesome has been highlighted as a key regulator in the control of anti-tumour immune responses in numerous solid cancer types ^23,24^. This key signalling hub consists of cell surface transmembrane integrin heterodimers which bind the extracellular matrix (ECM), as well as intracellular signalling adaptor proteins and kinases that control actomyosin machinery to mediate cancer cell behaviours such as invasion and migration ^24-27^. One of the major integrin adhesome signalling hubs is centred around focal adhesion kinase (FAK) which plays an essential role in regulating key cellular functions such as migration and cell adhesion ^24^, both in normal and malignant cells. Furthermore, FAK is known to control cell migration, invasion, metastasis and changes to growth and metabolism in glioblastoma models ^28-30^, with high expression of FAK being linked to lower survival in GBM patients ^28^. As such, FAK is considered to be a promising drug target for this disease ^31^.

FAK has strong immunomodulatory potential in some solid epithelial cancers that are receptive to immunotherapy, such as squamous cell carcinoma, non-small cell lung cancer and breast cancer, and in hard to treat, immunologically ‘cold’ solid tumours such as pancreatic ductal adenocarcinoma (PDAC) ^23,24,32-35^. Specifically, disruption of FAK’s function results in modulation of pro- and anti-inflammatory cytokines, impediment of pro-tumour stromal cells, increasing infiltration of anti-tumour immune cells, and overall boosting of anti-tumour T cell driven immune responses ^23,24^.

Understanding the regulators of the suppressive GBM tumour immune microenvironment may allow for greater success for immunotherapies in GBM treatment regimes. Here, we addressed whether FAK has key immunomodulatory roles within the GBM TME, and if so, how it functions. We considered whether FAK may operate both within GBM cells themselves, and/or in peripherally derived immune cells. We used genetic FAK deletion and a clinically relevant systemically delivered FAK inhibitor in a stem cell driven murine model of GBM. We identified a previously unrecognised mechanism whereby FAK controls peripherally derived monocyte trafficking to the GBM TME. We found that inhibition of FAK in monocytes causes eradication of suppressive macrophage populations within the tumour microenvironment and this is associated with reduced T cell dysfunction resulting in tumour clearance and significantly increased overall mouse survival.

## Results

### FAKi leads to GBM tumour clearance and increased survival

FAK inhibitors are currently being investigated in clinical trials for a wide range of solid tumour types ^23,24^, but have not yet been investigated for their potential immunomodulatory capacity in GBM. We chose to investigate the effects of FAK inhibition using a small molecule FAK inhibitor (FAKi) VS-4718, which has demonstrated target inhibition and efficacy in GBM murine models previously ^31^, and has been shown to modulate the anti-tumour immune response in other solid tumour types ^32,34^. We compared the immunomodulatory effects of tumour cell-intrinsic FAK gene deletion with FAKi. Tumours were established in mice following intracranial injection of the previously established 005-GSC GBM stem cell line ^36,37^, and treated with FAKi for 3 weeks at day 10 post tumour cell implantation at a concentration we have previously shown to inhibit FAK in GBM models ^31^. This resulted in a significant increase in survival when compared to control vehicle treated mice (Figure.1A), and we observed complete tumour clearance in 58.3% of FAKi treated mice, with 5/12 mice classified as non-responders (Figure.1B and Supplementary Figure.1A). FAKi treatment of a genetically distinct stem cell GBM model, namely mGB2 ^38^, also resulted in an increase in tumour free mice at day 200 (Supplementary Figure.1B), demonstrating its potency across models.

**Figure. 1.**
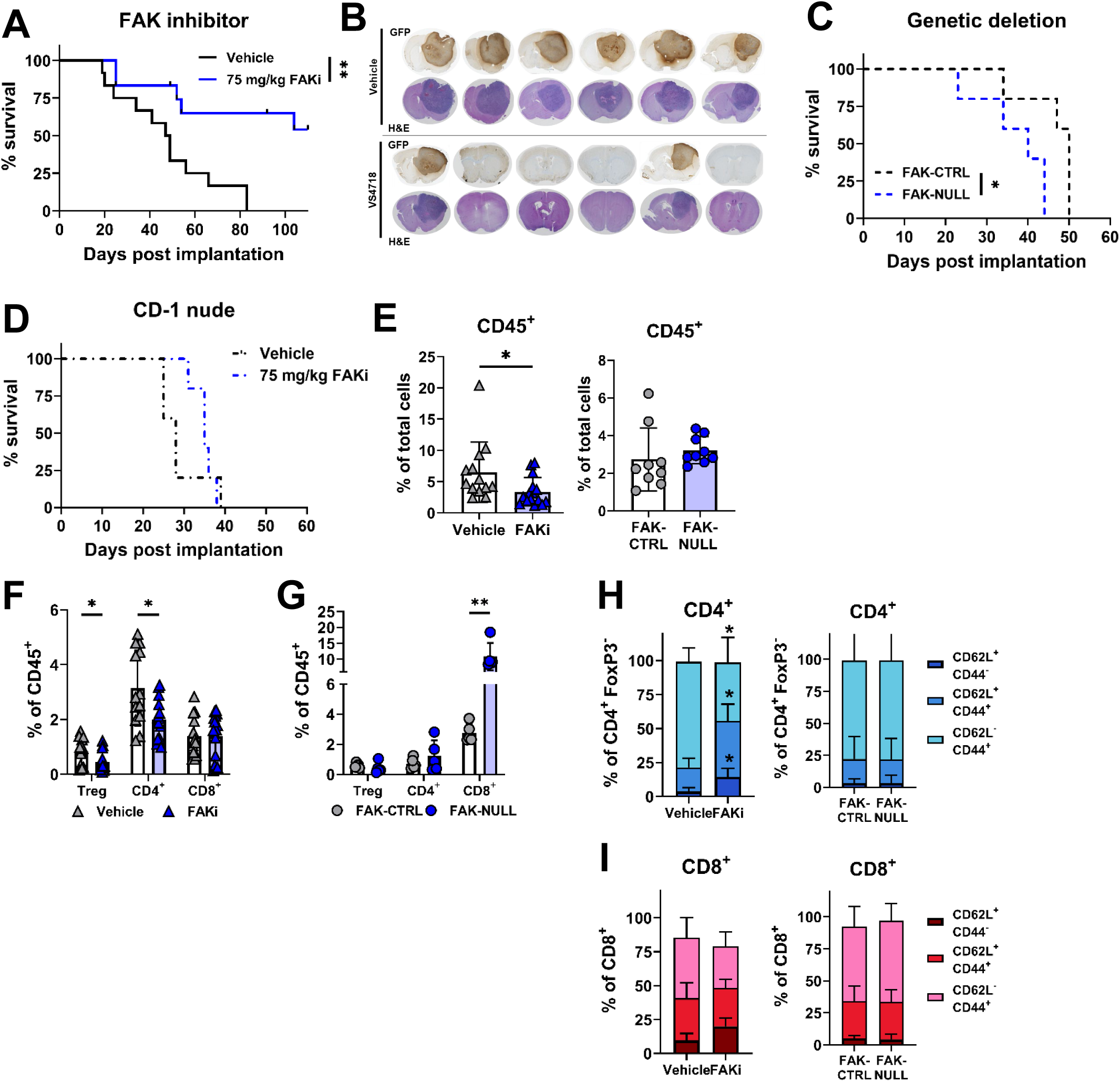
FAK inhibitor VS-4718 causes tumour clearance in a GBM model. **A)** C57BL/6 mice were intracranially implanted with 2x105 005-GSC cells, treated for 21 days with 75 mg/kg FAKi (VS-4718) or vehicle (HPMC) at day 10 post implantation, and monitored for overall survival. Combined data from 2 individual experiments, n=12 mice per group, statistical significance assess by Log Rank test. **B)** H&E and GFP immunohistochemistry of brains taken from one experiment in A. **C)** C57BL/6 mice were intracranially implanted as in A with either 005-GSC FAK-CTRL or FAK-NULL cells and monitored for overall survival. n=5 mice per group, statistical significance assessed by Log Rank test. **D)** As in A for CD-1 nude mice. n=5 mice per group, statistical analysis assessed by Log Rank test. **E-I)** As in A and C however mice were culled at Day 7 post FAKi treatment, or Day 17 post FAK-CTRL/NULL implantation. Tumour containing brain hemisphere was processed for immunophenotyping analysis of T cell populations. Gating strategy shown in Supplementary Figure.2A. Data shown: E) percent of CD45^+^ immune cells (as percentage of total live cells cells). F) Percent of T cell subsets (as percent of CD45^+^ cells) for FAKi treated tumours, and FAK-CTRL/NULL tumours (G). H) Expression of memory subset markers as a percent of total CD4^+^ FoxP3^+^ cells. with Naïve (CD62L^+^ CD44^-^), effector (CD62L^-^ CD44^+^) and central (CD62L^+^ CD44^+^) memory subsets. I) As in H as a percent of total CD8^+^. Each point = one mouse, representative examples shown of experiments performed at least twice, with statistical significance assess by unpaired t-tests. Significance denoted as p=<0.05*, p=<0.01**.

To determine whether tumour clearance was a function of inhibiting FAK within the tumour cells, we generated FAK gene deficient (FAK-NULL) 005-GSC cells and 005 CRISPR-Cas9 control (FAK-CTRL) cells (Supplementary Figure.1C). We found that loss of tumour cell-intrinsic FAK did not result in improved survival (Figure.1C) as we observed upon FAKi treatment, which was confirmed with a distinct second FAK-NULL CRIPSR-Cas9 cell line (Supplementary Figure.1D). Furthermore, neither genetic deletion of FAK (Supplementary Figure.1E) or direct inhibition with FAKi (Supplementary Figure.1F+G), impacted the size (area) of 005-GSC tumour spheroids grown *in vitro*. This demonstrates that the effect of FAKi treatment *in vivo* is not a direct effect on 005-GSC proliferation. Importantly, the increased survival benefit of FAKi treatment was ablated when the same treatment was given to immunodeficient CD-1 nude mice bearing 005-GSC tumours (Figure.1D), with no tumour clearance evident in this mouse strain that lacks adaptive immune system components, such as T cells. Together, these data demonstrated that loss of tumour intrinsic FAK expression is not sufficient to account for FAKi-induced GBM tumour clearance and requires the presence of a functioning adaptive immune system.

To investigate the impact of FAK modulation on anti-tumour immunity directly, tumours were removed at day 17 post tumour cell inoculation for both FAKi treated (day 7 post treatment initiation) and FAK-NULL tumour bearing mice, and adaptive immune cell populations were assessed using flow cytometry (Supplementary Figure.2A+B). Comparable sizes of brain fragments were processed to aid comparison between the two groups (Supplementary Figure.2C+D). We found that FAKi treatment resulted in a significant decrease in the overall number of immune cells (CD45^+^) within the brain fragment, which was not seen in the case of FAK-NULL tumours (Figure.1E). By contrast, a significant increase in total T cells (CD3^+^) as percent of total CD45^+^ population, was present only in FAK-NULL tumours and not in FAKi treated tumours (Supplementary figure.3A+B). Additionally, FAKi treated tumours displayed a decrease in immune-suppressive Treg cells (CD4^+^ FoxP3^+^) and non-Treg (CD4^+^ FoxP3^-^) cells (Figure.1F), whereas a significant increase in CD8^+^ T cell infiltration was present in FAK-NULL tumours (Figure.1G). The increase in CD8^+^ T cell infiltration was also confirmed by immunohistochemistry, which showed that T cells were distributed throughout the tumour mass and not present in surrounding normal brain (Supplementary Figure.3C+D). Together, these data demonstrate a disparity in immune infiltrates between systemic FAK inhibition and tumour cell-intrinsic FAK gene deletion. This resulted in a significant increase in the CD8:Treg ratio, particularly in FAK-NULL tumours (Supplementary Figure.3E+F). However, as no increase in tumour clearance in the FAK-NULL tumours was observed, it suggests that Treg cells may not be the major cell type conducting T cell suppression in this model. Therefore, changes in T cell subset numbers cannot account for the differences seen in survival outcomes between FAKi-treated and tumour cell-intrinsic FAK deletion.

As trafficking of immune cells to the TME is dependent on mature and functional vasculature ^39^, and FAK is known to directly regulate endothelial cell function ^40^, we addressed whether modulation of FAK in our GBM model resulted in changes to endothelial blood vessel architecture. To do this we implanted 005-GSC cells into Cdh5-CreER^T2^; Rosa26^lsl-tdT^ (tdTomato) reporter mice ^41,42^ in which the endothelial cells express tdTomato. There was no significant change in tdTomato^+^ blood vessel density following FAKi treatment (Supplementary Figure.3G+H), implying that FAKi treatment did not directly impact endothelial cell function.

CD8^+^ and CD4^+^ T cells infiltrating FAKi treated tumours and FAK-NULL tumours were quantified as naïve (CD44^-^ CD62L^+^), effector (CD44^+^ CD62L^-^) and central (CD44^+^ CD62L^+^) memory subsets (Figure.1H+I). FAKi treatment resulted in significant redistribution of all memory CD4^+^ T cell subsets (Figure.1H), with CD8^+^ T cell memory subsets unchanged (Figure.1I). This was likely driven by an increase in the proportion of naïve CD4^+^ T cells specifically (Supplementary Figure.3I). By contrast, this was not observed for FAK-NULL tumours, demonstrating that it was an effect that was not replicated by tumour cell intrinsic FAK inhibition (Figure.1H+I). These data suggest that systemic FAK inhibition leads to a reduction in the number of suppressive T cells and an increase in naïve CD4^+^ T cells in the TME, implying that FAKi treatment is capable of mobilising peripheral subsets of CD4^+^ cells that have the potential for tumour cell killing, potentially resulting in tumour clearance.

### FAKi leads to decreased effector T cell exhaustion

To understand the mechanism behind increased survival in FAKi treated GBM-bearing mice we examined changes in T cell phenotypes. FAKi treatment increased the expression of several key activation-associated markers (Figure.2A and individual mouse data shown in Supplementary Figure.4A+B) and there was a significant increase in CD107a expression and Granzyme B production in CD4^eff^ T cells, whereas this was not the case with CD8^eff^ T cells. Thus CD4^+^ T cells may have greater cytotoxic potential than CD8^+^ T cells, or are under less suppressive pressure within the TME of FAKi treated tumours. By comparison, CD8^eff^ and CD4^eff^ cells in FAK-NULL tumours had no changes in Granzyme B (Supplementary Figure.4C), indicating that depleting tumour cell FAK is not sufficient to impact T cell cytotoxicity and tumour cell killing, again highlighting the different effects of tumour cell FAK loss and systemic inhibition of FAK.

**Figure. 2.**
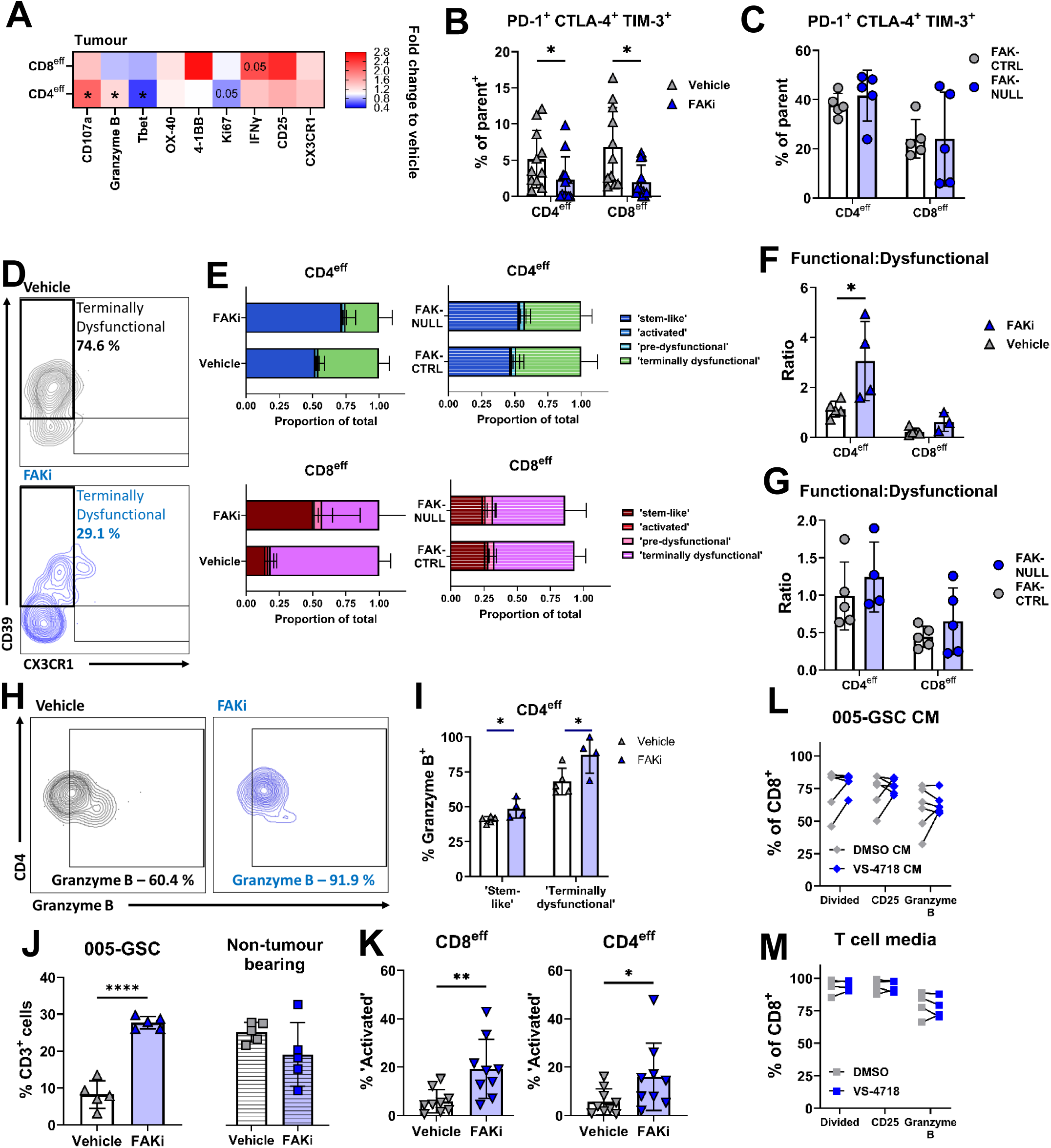
FAKi VS-4718 suppresses effector T cell exhaustion and induces GBM clearance. **A)** C57BL/6 mice were intracranially implanted with 005-GSC cells, treated for 7 days with 75 mg/kg VS-4718 or vehicle (HPMC) at day 10 post implantation, and mice were culled at Day 7 post VS-4718 treatment. Tumour containing brain hemisphere was processed for immunophenotyping analysis of T cell functional markers. Specifically: Heat map of markers associated with T cell activation and cytotoxic function shown as fold change over mean of vehicle, for CD4^eff^ (CD4^+^ FoxP3^-^ CD44^+^) and CD8^eff^ (CD8^+^ CD44^+^). **B)** Percent of CD4eff and CD8eff T cells triple positive for PD-1, CTLA-4 and TIM-3 as percent of parent population. **C)** C57BL/6 mice were intracranially implanted as in A with either 005-GSC FAK-CTRL or FAK-NULL cells, and tumours processed as in A. Percent of CD4^eff^ and CD8^eff^ T cells triple positive for PD-1, CTLA-4 and TIM-3 as percent of parent population. **D-G)** Assessment of T cell exhaustion subsets using gating strategy demonstrated in Supplementary Figure.4F, with ‘Stem-like’ (Ly108^+^ CX3CR1^-^), ‘Activated’ (Ly108^-^ CX3CR1^+^ CD39^-),^ ‘Pre-dysfunctional’ (Ly108^+^ CX3CR1^+^ CD39^-^) and ‘Terminally dysfunctional’ (Ly108^-^CX3CR1^-^ CD39^+^), with example of raw gating shown for terminally dysfunctional CD4^eff^ T cells shown. Populations shown as a proportion of total cells analysed (**E**). Ratio of functional subsets (stem-like+activated) to dysfunctional subsets (pre-dysfunctional+terminally dysfunctional) for FAKi treated tumours (**F**) and FAK-CTRL/NULL tumours (**G**). **H+I**) Expression of Granzyme B in ‘Stem-like’ and ‘Terminally dysfunctional’ populations as a percent of parent population in CD4^eff^, with example raw flow cytometry gating showing in **H**, quantification in **I**. **J)** Total T cells (CD3^+^) as a percent of total cell in whole blood collected from mice in A (**J** - left) and from non-tumour bearing mice treated with FAKi (**J** – right). **K)** Percentage ‘Activated’ subset as percent of CD8^eff^ (left) and CD4^eff^ (right) of whole blood T cells **L+M**) Flow cytometry analysis of percent divided, CD25 expression and granzyme B expression on in vitro stimulated T cells in presence of conditioned media from FAKi treated 005-GSC cells (**L**) or FAKi direct addition to T cells (**M**). Each point = one mouse, n=3-11 per group. Representative examples shown of experiments performed at least twice, with statistical significance assess by unpaired t-tests. Significance p=<0.05**\***, p=<0.01**\*\*,** p=<0.001*** and p=<0.0001****.

T cell exhaustion is widely regarded as a driving factor in allowing tumours to evade immune destruction ^43,44^. Therefore, we next investigated whether FAKi treatment modulated T cell exhaustion phenotypes. When 005-GSC tumour bearing mice were treated with FAKi, we observed a significant decrease in both CD4^eff^ and CD8^eff^ T cells that expressed key exhaustion markers PD-1 and CTLA-4, but not TIM-3 (Supplementary Figure.4D+E), and there was a significant reduction in T cells positive for all three receptors in the TME (Figure.2B) consistent with an increased tumour cell killing capability. In comparison, loss of tumour cell FAK after genetic deletion did not result in significant changes to effector T cell exhaustion markers (Figure.2C). In order to further investigate the functionality of the T cells, a more comprehensive investigation into T cell exhaustion was conducted using previously identified markers to define stages of T cell dysfunction^43^. We identified ‘stem-like’, ‘Activated’, ‘Pre-dysfunctional’ and ‘Terminally dysfunctional’ phenotypes within both CD8^eff^ and CD4^eff^ T cell populations (Supplementary Figure.4F). We also found a significant increase in ‘Stem-like’ CD4^eff^ cells within the GBM TME, with a corresponding significant decrease in ‘terminally-dysfunctional’ CD4^eff^ and CD8^eff^ cells (Figure.2D+E). As expected, this was associated with a significantly greater functional to dysfunctional ratio within the population of CD4^eff^ T cells (Figure.2F). By comparison, we did not observe significant changes in any of the exhaustion state T cell subsets in mice bearing FAK-NULL tumours (Figure.2E+G and Supplementary Figure.4G+H) Furthermore, ‘stem-like’ and ‘dysfunctional’ CD4^eff^, but not CD8^eff^ T cells were found to have increased granzyme B production in FAKi treated mice (Figure.2H+I and Supplementary Figure.4I), consistent with increased cytotoxic capacity in the CD4^eff^ T cell subset. Taken together, these data demonstrate that systemic FAK inhibition, but not tumour specific FAK deletion, can dampen terminal dysfunction in tumour infiltrating T cells and increase the cytotoxic potential of CD4^eff^ T cells.

We also examined the effects of FAKi treatment on circulating and peripheral T cells. There was a significant increase in total T cell number in the blood of FAKi treated tumour bearing mice (Figure.2J), but not in the spleen or bone marrow (Supplementary Figure.5A). Furthermore, this was not replicated in non-tumour bearing mice (Figure.2J), implying that the presence of a GBM tumour is altering mobilisation of T cells, as has been demonstrated previously for intracranial tumours ^45^. Although there was no proportional change in CD8^+^, CD4^+^ and Treg cells in the blood (Supplementary Figure.5B), a significant increase in ‘activated’ CD8^eff^ and CD4^eff^ cells was observed (Figure.2K), Together, these data demonstrate that systemically delivered FAKi induces mobilisation of activated peripherally circulating T cells, but only when a brain tumour is present.

Thus, our data demonstrate that FAKi treatment is capable of modulating the phenotype of tumour-infiltrating T cells, driving them away from terminal exhaustion and towards an increased cytotoxic state. As this was not evident following genetic deletion of FAK in the GBM cells themselves, it demonstrates that eradication of tumour cell intrinsic FAK expression is not sufficient to stimulate an effective anti-GBM T cell response and tumour clearance as seen following systemic treatment with the FAKi.

### Modulation of T cell phenotype is not a direct result of FAKi treatment

As tumour secreted factors are known modulators of immune infiltration and immune phenotypes within the TME ^46^, we assessed the impact of the tumour cell secretome on T cell function *in vitro* using 005-GSC conditioned media. Addition of conditioned medium from FAKi treated 005-GSC cells did not impair CD8^+^ T cell function or proliferation after anti-CD3 and anti-CD28 stimulation (Figure.2L), indicating that it is not inhibition of tumour cell intrinsic FAK that is causing changes to secreted factors which in turn can impact T cell activation.

We next asked whether the changes to T cell phenotypes seen after treatment with a systemic FAKi could be dependent on direct inhibition of FAK or its homologue Pyk2 that is expressed in T cells. T cells do not express FAK but do express Pyk2 ^47^, which is also targeted (albeit at lower potency) by the VS-4718 FAKi used *in vivo* ^48^. We assessed activation and cytotoxic potential of CD8^+^ T cells after exposure to VS-4718 *in vitro* at the same dose used above for FAK inhibition in the cancer cells (Supplementary Figure.1F). We found no significant difference in CD25 or granzyme B expression or in proliferative capacity when exposed to FAKi (Figure.2M). These data demonstrate that the FAK/Pyk2 inhibitory capacity of the FAKi does not directly impair T cell function and proliferation *in vitro*, indicating that the modulation of T cell exhaustion phenotypes seen with FAKi treatment is likely controlled by TME driven mechanisms.

### Systemic FAKi treatment modulates myeloid cell numbers and activation state

As the reduced T cell exhaustion, better tumour clearance and increased survival in FAKi treated mice was not caused by inhibition of tumour cell- or T cell-intrinsic FAK/Pyk2, this implies that different systemic mechanisms are involved. While T cells mediate the majority of tumour cell killing, it was important to assess the impact of FAK modulation on myeloid subsets within the TME. This is particularly relevant in GBM as myeloid infiltrates constitute the majority of immune cells within human GBM and play a significant role in immunosuppression of anti-tumour responses ^10,19,20^. Tumour bearing mice were treated with FAKi and alterations in myeloid lineages were assessed using flow cytometry. FAKi treatment led to a significant reduction in both peripherally-derived monocyte and macrophage (Mac) populations within the 005-GSC tumours, and an increase in the percentage of brain resident microglia (Figure.3A-E). No significant changes in neutrophils were observed, although dendritic cell (DC) percentages were lower in FAKi treated tumours (Supplementary Figure.6A+B). Importantly, this reduction in peripherally derived monocytes and Macs was only observed in FAKi treated tumours and not in the FAK-NULL tumours (Figure.3A-E), showing that systemic inhibition of FAK is required, as is the case for changes in the T cell compartment. Additionally, the reduction in monocytes/Macs requires the presence of tumour, as in non-tumour bearing mice who received FAKi there were no differences in monocytes or Mac populations within the brain (Supplementary Figure.6C).

**Figure. 3.**
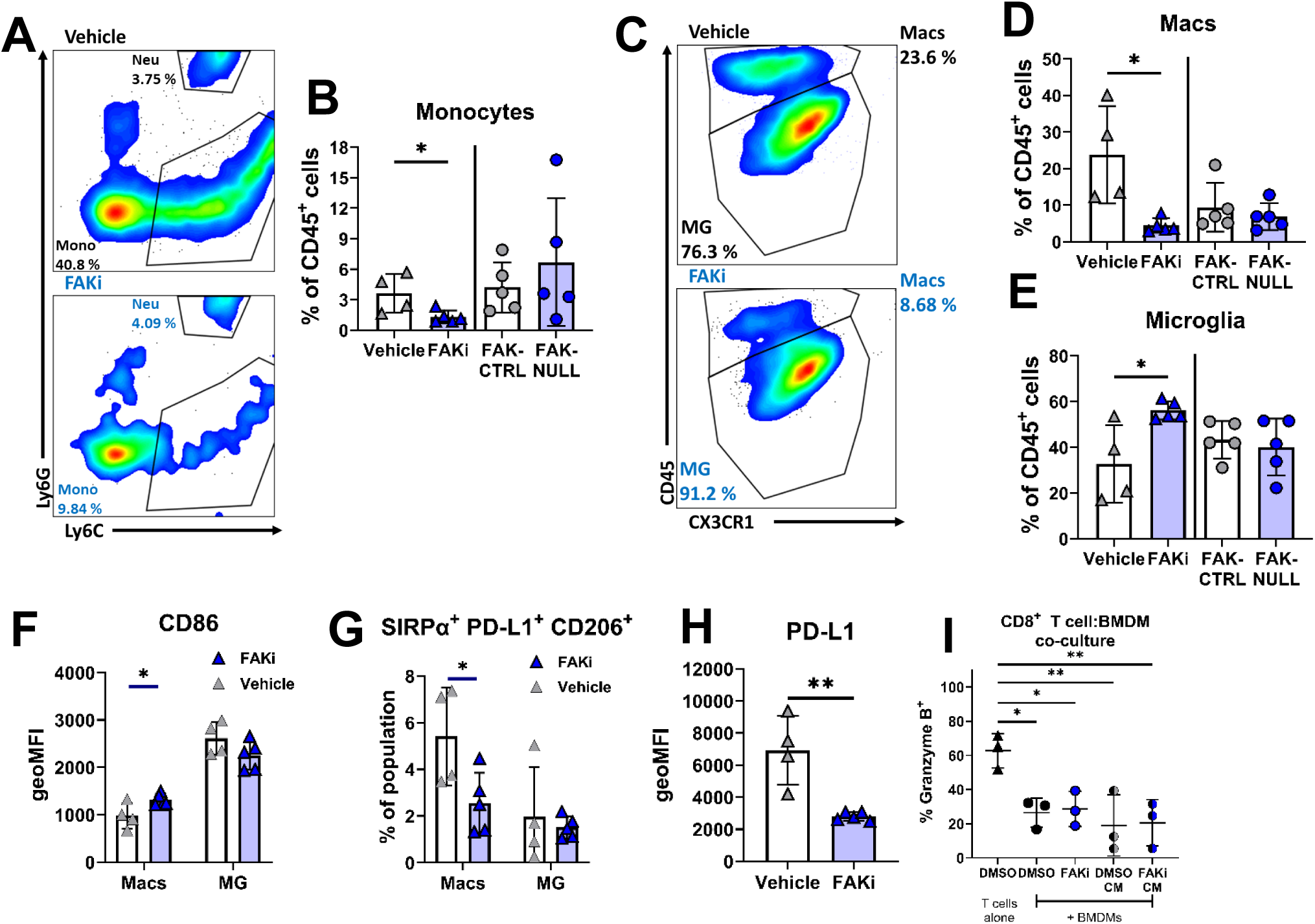
FAKi VS-4718 but not tumour cell FAK deletion suppresses immunosuppressive peripherally derived monocyte populations. **A-E**) C57BL/6 mice were intracranially implanted with 005-GSC cells, treated for 7 days with 75 mg/kg FAKi (VS-4718) or vehicle (HPMC) at day 10 post implantation, and mice were culled at Day 7 post treatment. Alternatively, C57BL/6 mice were intracranially implanted as in A with either 005-GSC FAK-CTRL or FAK-NULL cells, and mice were culled at Day 17. Tumour containing brain hemisphere and bone marrow was processed for immunophenotyping analysis of myeloid populations, gating shown in Supplementary Figure.2B. Specifically: Percentage of tumour infiltrating monocytes (**A+B**), Macrophages (**C+D**), Microglia (**C+E**), as a percent of total immune cells (CD45^+^). Each point = one mouse, representative examples shown of experiments performed at least twice, with statistical significance assess by unpaired t-tests. **F-H**) As in **A**. Expression as mean fluorescence intensity (geoMFI) of CD86 on both Macrophages (Macs) microglia (MG) (**F**). Percentage of Macs and MG triple positive suppressive associated markers SIRPα, PD-L1 and CD206 shown as a percent of parent population (**G**). Expression as geoMFI of PD-L1 on tumour infiltrating Macs (**H**). Each point = one mouse, representative examples shown of experiments performed at least twice, with statistical significance assess by unpaired t-tests. **I)** Percent of CD8^+^ T cells expressing granzyme B in in vitro co-cultures with BMDMs with FAKi treatment or in the presence of conditioned media from 005-GSC treated with FAKi. Significance p=<0.05**\***, p=<0.01**\*\***.

As myeloid cells, and particularly Macs, are regarded as major contributors to immunosuppression in GBM ^10,19^, it is important to understand how these infiltrating Macs are polarised when exposed to FAKi treatment. To that end, phenotypes of myeloid cells were assessed using an array of markers of ‘activated’ (CD86) and ‘suppressive’ (CD206, SIRPα and PD-L1) macrophage polarisation ^49-52^. Of note, Macs, but not microglia (Figure.3F), had significantly increased CD86 expression after FAKi treatment, suggesting that they were more capable of co-stimulating T cell responses, and so more ‘activated’ ^52^. Repolarisation of Macs away from a ‘suppressive’ state following FAKi treatment was confirmed with a significant reduction in Macs expressing SIRPα, PD-L1 and CD206, with no significant changes to microglia (Figure.3G). This was mainly driven by a significant downregulation of PD-L1 expression on Macs within the TME of mice treated with FAKi (Figure.3H), with no changes in PD-L1 expression on other myeloid subsets examined (Supplementary Figure.6D). In addition, changes in PD-L1 expression were not observed in tumour Macs from FAK-NULL tumour bearing mice (Supplementary Figure.6E). These data show that FAKi treatment resulted in the reduced presence of suppressive Macs within the GBM TME.

To understand whether the FAKi was directly impacting the suppressive capacity of macrophages, co-culture assays of bone marrow derived macrophages (BMDMs) with CD8^+^ T cells in the presence of FAKi were conducted *in vitro* (Figure.3I). Here, addition of BMDMs to T cells significantly decreased their Granzyme B production compared to T cells alone. However, neither direct addition of FAKi to the co-culture, nor addition of conditioned media from FAKi treated 005-GSC cells significantly altered T cell Granzyme B production compared to vehicle treated BMDM cultures, demonstrating that secreted factors from FAKi treated tumour cells are not directly impacting macrophage suppressive capacity. Furthermore, key macrophage phenotype markers, PD-L1 and MHC II, were unaltered by the direct addition of FAKi *in vitro* (Supplementary Figure.6F-G). However, addition of conditioned media from FAKi treated 005-GSC cells did result in an increase in CD206 (Supplementary Figure.6H).

### Systemic FAKi treatment leads to sequestering of inflammatory monocytes in the bone marrow

We next used flow cytometry to assess how the FAKi effects peripheral reservoirs of monocytes in 005-GSC tumour bearing mice. We did not observe any changes in splenic monocytes or circulating monocytes in the blood, but did find a significant increase in monocytes within the bone marrow (Figure.4A), while other myeloid subsets were unchanged (Supplementary Figure.7A+B). This was dependent on the presence of a tumour, as non-tumour bearing mice treated with FAKi did not display the same increase in bone marrow monocytes, but was not dependent on tumour cell-intrinsic FAK loss (Figure.4A). FAK inhibition led to an increase in the ratio of inflammatory (Ly6C^hi^ CX3CR1^+^ CCR2^+^) to patrolling (Ly6C^lo/int^ CX3CR1^lo^ CCR2^-^) monocytes, demonstrating an accumulation of inflammatory monocytes within the bone marrow (Figure.4B). Therefore, the systemic effects of FAKi treatment were driving an increase in inflammatory bone marrow monocytes and a concurrent decrease in Macs within tumours.

**Figure. 4.**
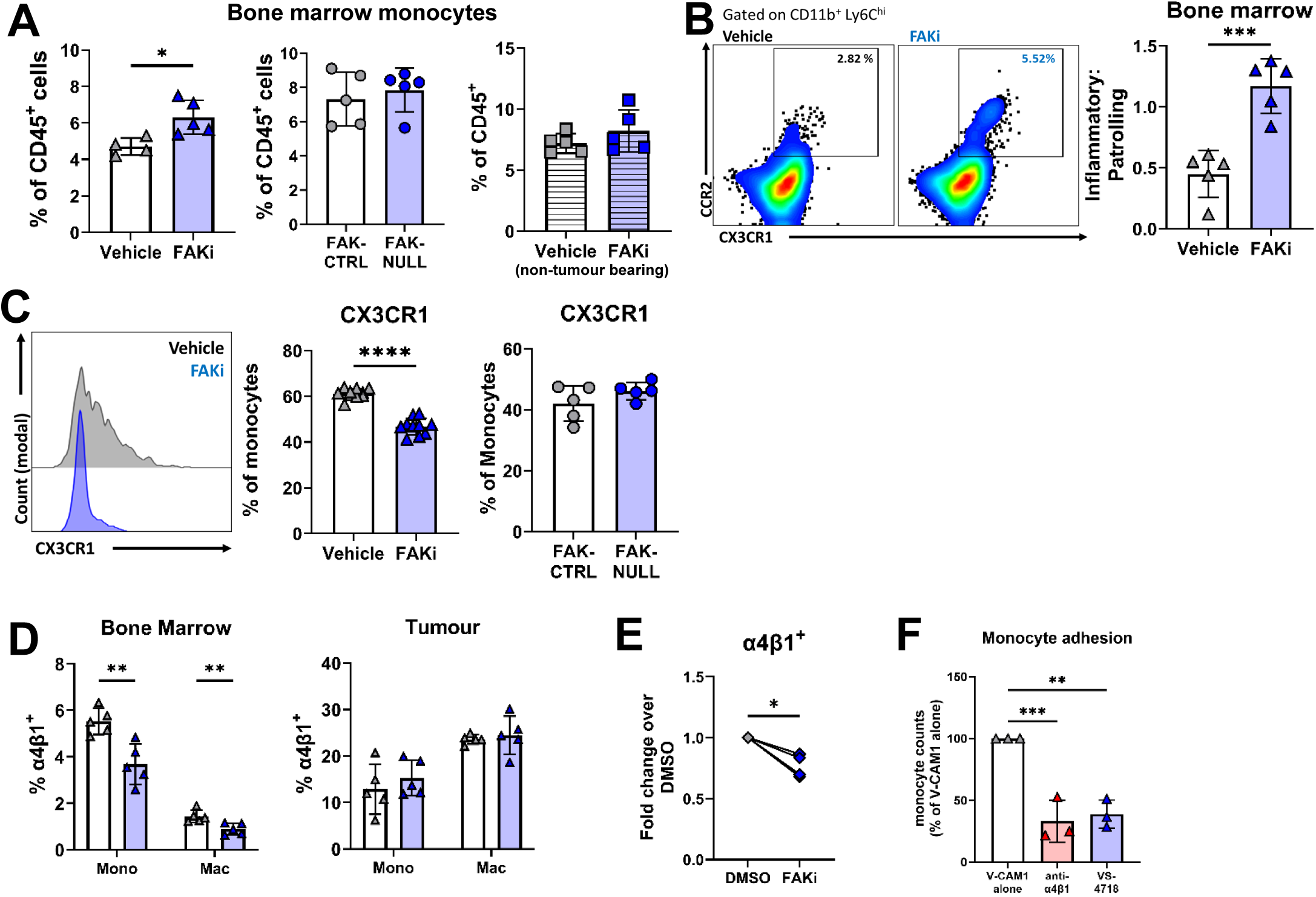
FAKi VS-4718 promotes sequestration of inflammatory monocytes in the bone marrow and suppression of integrin dependent adhesion. **A)** C57BL/6 mice were intracranially implanted with 005-GSC cells (or sham surgery performed for non-tumour bearing), treated for 7 days with 75 mg/kg VS-4718 or vehicle (HPMC) at day 10 post implantation, and mice were culled at Day 7 post VS-4718 treatment. Alternatively, C57BL/6 mice were intracranially implanted as in A with either 005-GSC FAK-CTRL or FAK-NULL cells, and mice were culled at Day 17. Tumour containing brain hemisphere and bone marrow was processed for immunophenotyping analysis of myeloid populations and functional markers, gating of populations shown in Supplementary Figure.2B. Specifically: bone marrow monocytes as a percentage of total immune (CD45^+^) cells in FAKi treated mice (left), FAK-NULL tumour bearing mice (middle) and non-tumour bearing mice (right). **B)** As in A with (**B** – left) of ‘inflammatory’ (Ly6C^hi^ CX3CR1^+^ CCR2^+^) and ‘Patrolling’ (Ly6C^lo/int^ CX3CR1^lo^ CCR2^-^) monocytes in bone marrow, with quantification as a ratio of ‘inflammatory’ to ‘patrolling’ monocytes (**B** – right). **C)** Histograms of CX3CR1 expression on total (CD11b^+^ Ly6C^+^ Ly6G^-^) bone marrow monocytes (**C** – left), with quantification of number of monocytes CX3CR1^+^ as a percent of parent population in tumour bearing mice (**C** – middle), and FAK-NULL tumour bearing mice (**C** – right). **D)** Percent of bone marrow (**E** – left) and tumour (**E** – right) monocytes and macrophages positive for α4β1 (VLA-4) integrin as a percent of parent populations. **E)** Expression of α4β1 on in vitro isolated monocytes after FAKi treatment, as a fold change over DMSO control. **F)** VCAM-1 adhesion assay of monocytes in the presence of FAKi, with anti-α4β1 antibody as positive control. Each point = one mouse, representative examples shown of experiments performed at least twice, with statistical significance assess by unpaired t-tests. Significance p=<0.05**\***, p=<0.01**\*\*,** p=<0.001*** and p=<0.0001****.

The change in abundance of monocytes we observed within the bone marrow was likely not due to the ability of the FAKi to directly alter the proliferation of monocytes, as there was no change in Ki67 expression in the TME and bone marrow monocyte populations (Supplementary Figure.7C+D). This suggests that the increased numbers of monocytes in the bone marrow and reduced TME myeloid populations may be due to their inability to egress into the circulation and extravasate within the TME.

To address how FAKi treatment leads to a reduction in the presence of suppressive Macs within the TME, we first examined the CCL2/CCR2 axis. The cytokine CCL2, via binding to its cognate receptor CCR2, is a major driver of bone marrow derived monocyte recruitment to sites of inflammation ^53,54^, and tumours, including GBM ^55-57^. We found no changes in CCR2 expression levels on monocytes or Macs in peripheral tissues reservoirs following FAKi treatment (Supplementary Figure.8A-C); likewise, there were no changes in CCL2 production by 005-GSCs treated with FAKi *in vitro* (Supplementary Figure.8D).

We next addressed whether tumour cell-intrinsic deletion of the gene encoding FAK altered the secretion of other cytokines that are involved in recruitment and migration of myeloid cells. There was no difference in CXCL1, CXCL4 or CCL12 secretion into the conditioned medium collected from FAK-CTRL and FAK-NULL cells. There was a modest, but statistically significant, increase in IL-6 production (Supplementary Figure.8E+F).

Previous studies have shown that loss of FAK in monocytes/macrophages downregulates their adhesion and migration^58-60^. These processes require both chemokine receptors such as CX3CR1, and integrin α4β1 (VLA-4 – CD49d and CD29 respectively) which regulate the trafficking of monocytes to sites of inflammation and tumours ^61^. We found that FAKi treatment reduced the percentage of CX3CR1^+^ bone marrow monocytes (Figure.4C), but there was no reduction in CX3CR1^+^ bone marrow monocytes in FAK-NULL tumour bearing mice (Figure.4C). FAKi treatment also downregulated the expression of α4β1 on bone marrow monocytes, but not on those in the TME (Figure.4D+E). Additionally, no significant changes were seen in another important monocyte adhesion integrin heterodimer, α5β1 (VLA-5 – CD49e CD29 respectively) on the same cells (Supplementary Figure.8G), indicating that FAKi resulted in specific loss of α4β1. α4β1 dependent adhesion of monocytes to VCAM-1 on endothelial cells is required for their transendothelial migration and is a critical step in monocyte trafficking from the periphery to the sites of inflammation, such as the TME ^53^. Consistent with this, FAKi treatment of isolated bone marrow derived monocytes *in vitro* led to a significant reduction in α4β1 expression on monocytes and a reduction in adhesion to VCAM-1 (Figure.4E+F). While FAK regulates expression of VCAM-1 on endothelial cells that can also impact on transendothelial migration^62^, we found that VCAM-1 expressed on endothelial cells in the GBM TME and bone marrow following FAKi treatment was not significantly different (Supplementary Figure.8H+I).

Overall, these data demonstrate that systemically delivered FAKi can impair monocyte trafficking from the bone marrow and is associated with reduced expression of key monocyte adhesion molecules, CX3CR1 and α4β1.

### Impaired trafficking of FAK-deficient monocytes promotes increased T cell activation

To confirm that peripheral bone marrow derived monocyte lineages are present within the 005-GSC tumours, we utilised *Ms4a3*^Cre^:Rosa26^lsl-tdT^ mice ^63^, where cells of bone marrow derived granulocyte-monocyte (GMP) lineages, including monocyte derived macrophages (MDMs) are labelled with TdTomato. 005-GSC cells were implanted into brains of *Ms4a3*^Cre^:Rosa26^lsl-tdT^ mice and imaged at day 20 post implantation (Supplementary Figure.9A). As expected, we observed numerous bone marrow derived monocytes/MDMs present within the 005-GSC tumours, with few TdTomato^+^ cells seen outside the tumour mass (Supplementary Figure.9A+B). To ascertain that inhibition of FAK in monocytes was impairing their ability to migrate to the GBM TME, we created an adoptive transfer model using traceable *Ms4a3*^Cre^:Rosa26^lsl-tdT^ monocytes, which had FAK genetically deleted (sgFAK). sgFAK or sgCTRL monocytes were adoptively transferred intravenously into 005-GSC tumour bearing mice at day 15 post tumour implantation (Figure.5A+B and Supplementary Figure.9C). Tracking of TdTomato^+^ monocytes by flow cytometry was conducted 4 days later to assess changes to monocyte trafficking (Supplementary Figure.9D). When tracing TdTomato^+^ cells across tissues, we observed a significant reduction in total TdTomato^+^ cells in the TME of mice who received sgFAK monocytes, compared to sgCTRL monocytes (Figure.5C), resulting in a significantly lower ratio of TdTomato^+^ MDMs to native MDMs (Figure.5D). All other tissues assessed had low levels of TdTomato^+^ cells (Figure.5C), which was not impacted by FAK deletion. The majority of TdTomato^+^ cells within the TME had differentiated into macrophages (F4/80^+^ Ly6C^-^), of which there were comparable proportions of monocytes and macrophages in both sgFAK and sgCTRL monocyte treated groups (Figure.7E). In contrast, loss of FAK protein in monocytes via gene deletion did not significantly impact the proportions of inflammatory or patrolling TdTomato^+^ monocytes within the tumour (Supplementary Figure.9E). Importantly, the non-monocyte derived neutrophils and microglia displayed negligible TdTomato expression, confirming they were not derived from peripheral monocyte precursors (Supplementary Figure.9F). However, a small percentage of DCs were found to express TdTomato, which would account for monocyte-derived DCs ^53,64^. These data demonstrate that loss of FAK expression in peripheral monocytes impairs their ability to traffic to the GBM TME from the blood, but does not affect their ability to differentiate into macrophages within the tissue. Finally, sgFAK monocyte recipient mice displayed increased CD3^+^ T cell infiltration into the TME, correlating with the percentage of TdTomato^+^ cells present in the tumour (Figure.5F+G). CD8^+^ T cells within the GBM TME had increased Granzyme B production in mice receiving sgFAK monocytes (Figure.5H), indicative of increased cytotoxic potential. This demonstrates that the FAK dependent impairment of monocyte trafficking to the GBM TME, results in an increase in T cells with cytotoxic potential.

**Figure. 5.**
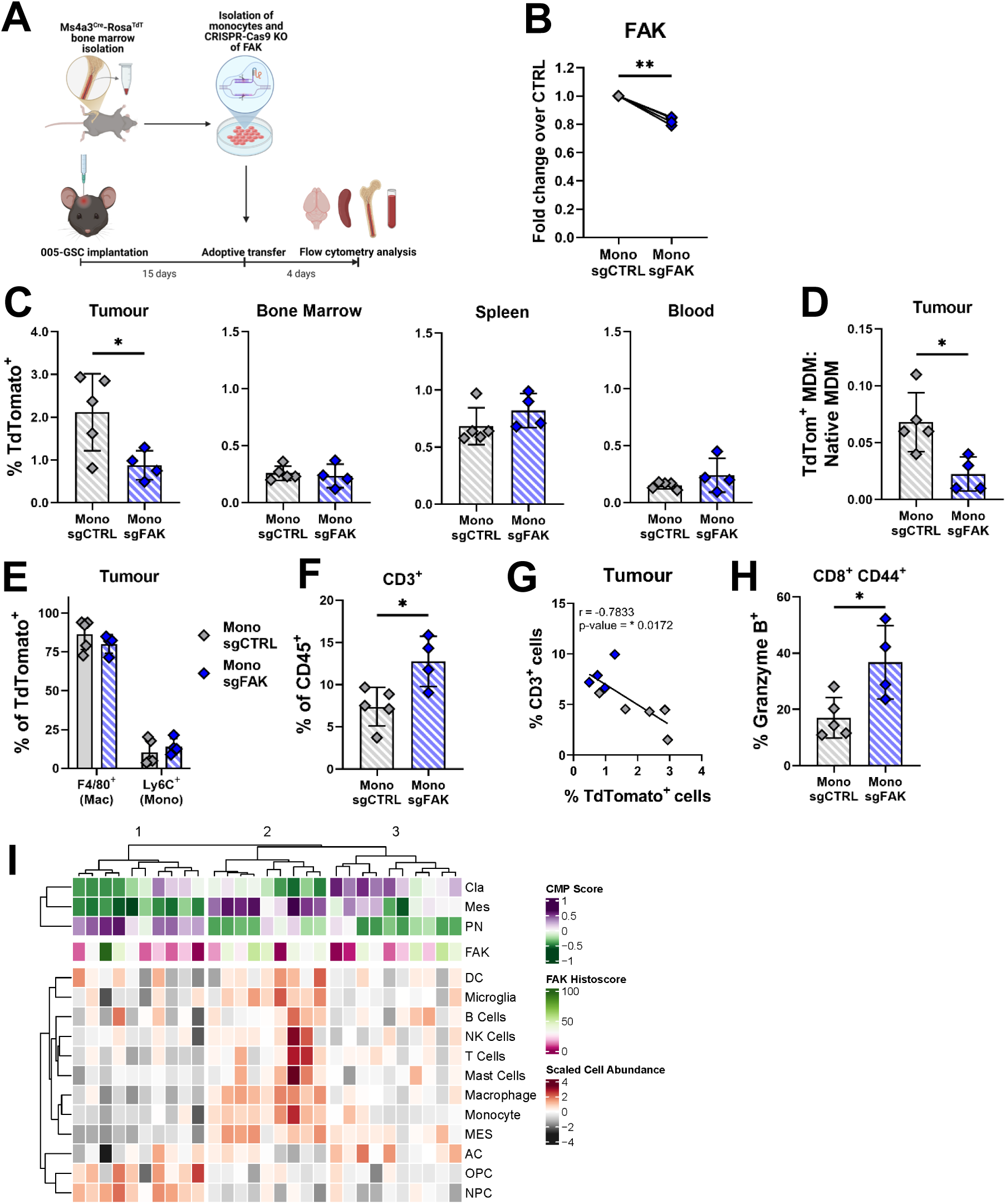
Monocyte cell-specific FAK gene deletion results in impaired trafficking to GBM tumours and correlates with increased T cell cytotoxic potential. **A)** Experimental schematic of adoptive transfer of FAK-deleted monocytes into 005-GSC tumour bearing mice. Male Ms4a3^Cre^-Rosa^TdT^ mice were used for monocyte collection, and were adoptively transferred into male C57BL/6 recipient mice. **B)** Quantification of FAK deletion in monocytes in vitro with example flow cytometry data shown in Supplementary Figure.9C **C)** Percent of total CD45^+^ cells expressing TdTomato in the tumour and systemic tissues in mice receiving FAK-CTRL or FAK-NULL monocytes. **D)** Ratio of TdTomato^+^ MDMs to naïve (TdTomato^-^) MDMs in tumour infiltrating MDMs **E)** Percentage of macrophages or monocytes which are TdTomato^+^ in 005-GSC tumours **F)** Numbers of total T cells (CD3^+^) as a percent of total CD45^+^ cells infiltrating into 005-GSC tumours of mice receiving either FAK-CTRL or FAK-NULL monocytes. **G)** Correlation between % of CD3^+^ cells and % of TdTomato cells in 005-GSC tumours **H)** Percentage of CD8^eff^ cells expressing Granzyme B in into 005-GSC tumours of mice receiving either FAK-CTRL or FAK-NULL monocytes. **I)** Heatmap showing classical (Cla), proneural (PN), and mesenchymal (Mes) bulk RNA scores for 29 patient tumours, matching FAK protein histoscores, and inferred deconvolved cell type abundances. Immune cells: B cells, DC – Dendritic cells, Macrophage, Mast cells, Microglia, Monocyte, NK cells, T cells. Neoplastic cells: AC – astrocyte-like, MES – mesenchymal cells, NPC – neuronal progenitor-like, OPC – oligodendrocyte progenitor-like. For adoptive transfer experiments: Each point = one mouse, with statistical significance assess by unpaired t-tests. Significance p=<0.05**\***, p=<0.01**\*\*,** p=<0.001*** and p=<0.0001****.

To understand whether expression of FAK in human GBM is correlated with changes in immune infiltration we examined both total FAK and phospho Y576-FAK (pFAK) expression across a previously detailed brain tumour tissue microarray (TMA) ^65^. 27 GBM tumours were evaluable, which revealed both nuclear and cytoplasmic FAK staining, with 11 tumours positive for pFAK staining (Supplementary Table.1). Interestingly, neither FAK histoscore or pFAK positivity correlated with percent of cells positive for macrophages markers CD163 or CD68 (Supplementary Table.2 and 3). This suggests that tumour intrinsic FAK expression does not impact macrophage infiltration, consistent with what was observed in the 005-GSC FAK-NULL mouse model. To confirm whether FAK expression impacts infiltration of different immune subsets, matched RNAseq data from the patients represented on the TMA was used. Here, samples were stratified into the previously defined transcriptional subtypes: classical (Cla), proneural (PN) and mesenchymal (Mes), based on gene expression ^66^. To explore whether tumour FAK is associated with immune cell abundance in patient tumours, we used a previously published deconvolution method ^67^ to infer immune and tumour cell type abundances and correlated these with tumour FAK. Notably, tumours with a higher Mes score were demonstrated to have the most abundant immune cell infiltration, compared to tumours with higher Cla or PN scores (Figure.5I). It has been previously documented that mesenchymal-like GBM tumours have higher levels of immune cell infiltration ^7,8,68^, and this was confirmed here with applying the same Scaled Cell Abundance scoring to a publicly available Clinical Proteomic Tumor Analysis Consortium (CPTAC) dataset ^69^ (Supplementary Figure.10A) of GBM patients, demonstrating that the relatively small TMA used here is transcriptionally representative of GBM. Importantly, FAK histoscore was not enriched in any of the transcriptional subtypes and did not significantly correlate with any of the immune cell types investigated (Figure.5I and Supplementary Figure.10B). Overall, in keeping with the murine model data, these data suggest that tumour intrinsic FAK expression in human GBM does not control infiltration of immune cells, and importantly monocytes/macrophages, into the TME. This indicates that a systemically acting FAKi is needed to modulate immune cell infiltration in patients.

## Discussion

GBM is characterised by a distinct lack of cytotoxic T cell infiltration and an abundance of peripherally derived immunosuppressive myeloid cells, specifically macrophages and their monocyte precursors, leading to a highly immunosuppressive TME ^10-12^. This significantly impacts on the ability of GBM to instigate an anti-tumour immune response, and it is therefore essential to establish strategies to generate a more anti-tumour conducive TME. The immune modulatory capacity of targeting FAK and other integrin adhesome proteins has been reported in a number of solid tumour types ^23,24^, yet FAKs role in intracranial tumours, and GBM specifically, is not known. Here, we report that while deletion of the gene encoding FAK in GBM cells resulted in an increase in T cell infiltration into the tumours grown in the brains of recipient mice, this did not result in an increase in overall mouse survival. Therefore, increasing T cell numbers alone is not sufficient to induce anti-tumour immunity. This is in contrast to our previous work in squamous cell carcinoma ^34^ and as reported in PDAC ^32^, where genetic deletion of FAK in the tumour cells was sufficient to impact anti-tumour immunity and increase survival. The lack of response in GBM, is most likely due to the highly heterogenous and myeloid cell rich environment, which is still capable of supressing anti-tumour T cell responses even with a greater influx of T cells.

Despite the lack of survival advantage following GBM cell specific FAK deletion, a systemically delivered FAKi significantly improved mouse survival and tumour clearance, which was reliant on the presence of adaptive immunity. Strikingly, systemic FAKi, and not GBM specific FAK gene deletion, led to less exhausted T cells with increased cytotoxic potential, particularly in the CD4^+^ (non-Treg) subset. Importantly, CD4^+^ T cells were recently shown to be crucial to the anti-tumour immune response in GBM ^70^, suggesting that CD4^+^ T cells have potential to be the drivers of tumour clearance in GBM models. Further investigation revealed that the reduction in T cell exhaustion after FAKi treatment was likely due to a significant decline in the presence of suppressive MDMs that are known to be important drivers of immunosuppression and T cell exhaustion in GBM ^44,70,71^. Previous studies have focussed on tumour cell intrinsic effects of FAK in modulating the TME, and indeed tumour intrinsic deletion of FAK in 005-GSCs was sufficient to increase the number of infiltrating CD8^+^ T cells and reduce the number of Tregs as has been described in other tumour types ^32,34^. We recognise that when treating with small molecule inhibitors, including those that target FAK kinase activity, it is not possible to rule out effects on multiple cell types; indeed, we found that inhibition of FAK in peripheral myeloid populations can impact their intratumoural recruitment. Interestingly, treatment with the same FAKi in murine models of PDAC, did not result in changes in numbers of monocytes within the bone marrow ^32^, suggesting here we are observing a GBM specific response. Furthermore, in a human GBM TMA cohort we demonstrated that tumour intrinsic FAK expression was not correlated with immune cell infiltration, demonstrating for the first time that tumour intrinsic FAK does not regulate immune cell recruitment in GBM. Both FAK (and its homolog Pyk2) are expressed in myeloid cells and although we found that specific deletion of FAK in monocytes was sufficient to suppress their recruitment and increase the number of cytotoxic T cells in the GBM tumours, we cannot rule out the formal possibility that Pyk2 may also contribute since the FAKi used in this study (VS-4718) also inhibits Pyk2 (although at significantly lower potency) ^48^. FAK and Pyk2 are both known to play a role in the motility of myeloid cells ^58-60^, with Pyk2 demonstrated to influence monocyte to macrophage differentiation in PDAC ^72^. However, this was shown to be dependent on TME stiffness, which will be different when comparing extracranial to intracranial tumours due to disparity in extracellular matrix composition ^73,74^. Furthermore, we demonstrated in the context of GBM, monocytes lacking FAK were not impaired in their capability to differentiate into MDMs, suggesting that may be a FAK independent mechanism.

Myeloid cell specific deletion of the gene encoding FAK reduced the recruitment of macrophages to sites of inflammation, indicating a broader role for FAK in the trafficking of myeloid cells ^58^. This was linked with FAK-dependent alterations in macrophage adhesion dynamics, chemotaxis and migration, and can be attributed to the pivotal role that FAK plays in mediating integrin-dependent adhesion. Consistent with this, we found that treatment with the FAKi in monocytes reduced their α4β1 integrin-dependent adhesion to VCAM-1. The FAK inhibitor also reduced levels of α4β1 on monocytes, showing that FAK can have additional effects on this integrin signalling pathway that is known to have a key role in monocyte trafficking ^53^. The mechanism(s) behind this are not known, but FAK modulates cytokine signalling pathways that regulate expression of α4β1 ^75-77^. The FAKi also suppressed the levels of the cytokine receptor CX3CR1 on monocytes. CX3CR1 together, with its ligand CX3CL1, regulate the adhesion of monocytes to endothelial cells and is another important pathway involved in monocyte trafficking ^53^. CX3CL1 can also bind and activate α4β1 ^78,79^, and since α4β1-VCAM-1 mediated adhesion is involved in the transendothelial migration of monocytes into tumours ^80^, this suggests one way in which FAK may be regulating the role of monocytes in GBM.

It is important to note that apart from the reduction in CX3CR1 expression, the majority of FAKi -induced changes on monocytes were only observed in tumour bearing mice and not in the sham surgery mice that bore no tumours. This highlights the systemic-wide impact that tumours, and GBMs in particular, can have on distant tissues such as bone marrow ^45,81^; in keeping with this, we found that treatment of BMDMs with 005-GSC conditioned medium significantly increased their ability to suppress the activity of CD8^+^ T cells. It is therefore important to consider changes in the organism as a whole, before, during and after immunomodulatory treatments such as with FAK inhibitors that are in clinical development, which can have both tumour cell intrinsic effects, and effects on immune cells at peripheral sites like the bone marrow.

We show here a completely novel role for systemic inhibition of FAK in peripheral monocyte cell populations that leads to an impairment in trafficking to the GBM tumours. This is likely due to a loss of key adhesion and migration signalling through α4β1 and CX3CR1 in peripheral monocytes. As a result, there are fewer monocytes in the TME and a consequent reduction in suppressive MDMs, a reduction in cytotoxic T cell exhaustion and enhanced GBM clearance and mouse survival. Our work here describes for the first time that targeting integrin-associated proteins on peripheral immune cell populations may be a useful strategy, and highlights a new mechanism of action by which systemic FAK inhibitors can act as beneficial immune modulatory therapies in GBM.

## Methods

### Cell lines

005-GSC (glioblastoma stem cells) cells were kindly provided by Dr Samuel Rabkin (Harvard Medical School, Department of Neurosurgery, Boston, MA, USA). The cells were originally developed as described previously ^37^, and have activated H-Ras, activated AKT on a p53^+/-^background. Cells were maintained at 37°C, 5% CO_2_ in Advanced Dulbecco Modified Eagle Medium (DMEM)/F12 reduced serum media (Gibco, #12634010) supplemented with 20 ng/ml EGF (Gibco, #315-09-1MG), 20 ng/ml FGF (Gibco, #450-33-1MG), 1% N2 supplement (Gibco, #17502048), 2 mM L-glutamine (Gibco, #25030081), 2 ug/ml heparin (Sigma-Aldrich, #H3393) and 0.5% penicillin-streptomycin (PS) (Gibco, #15140122). mGB2 cells were kindly provided by Prof. P. Angel, German Cancer Research Centre (Heidelberg, Germany). The cells were developed as described previously ^38^, and, in short, were generated from tumours derived from Tlx-CreERT2/p53^fl/fl^Pten^fl/fl^C57BL/6 mice by serial implantation into wildtype C57BL/6 mice over two rounds. Cells were maintained at 37°C, 5% CO_2_ in Dulbecco DMEM/F12 media (Gibco, #21331020) supplemented with 20 ng/ml EGF (Gibco, #315-09-1MG), 20 ng/ml FGF (Gibco, #450-33-1MG), 1% N2 supplement (Gibco, #17502048), 2 mM L-glutamine (Gibco, #25030081) and 1% PS (Gibco, #15140122).

### Mice

All experiments were carried out in accordance with UK Home Office regulations. All regulated procedures carried out were approved by the local Animal Ethics and Welfare Review Board, and performed in accordance with the Animals (Scientific Procedures) Acts 1986 as set out in project license: PP7510272. C57BL/6J and CD-1 nude (Crl:CD1-Foxn1^nu^) were purchased from Charles River, maintained by the Biomedical Research Facility staff at the University of Edinburgh and used between 7- 26 weeks of age. Ms4a3^Cre^-Rosa^TdT^ mice were originally generated by Liu et al. by backcrossing Ms4a3^Cre^ mice onto a C57BL/6 background and subsequently crossing with Rosa26TdTomato reporter mice ^63^, and were locally bred and maintained by the Biomedical Research facility at the University of Edinburgh. Cdh5-CreER^T2^; Rosa26^lsl-tdT^ (tdTomato) were created with backcrossing Cdh5-CreER^T2^ mice with Ai14 reporter mice ^41,42^. To induce red fluorescent VE-Cadherin (Cdh5), Cdh5-CreER^T2^; Rosa26^lsl-tdT^ mice were treated with Tamoxifen (100 mg/kg) by oral gavage daily for 5 days prior to cell implantation ^82^. All mice were female unless otherwise stated in figure legends. No blinding was performed.

### Intracranial implantation of 005-GSC tumour cells

005-GSC cell suspensions of 1×10^4^/ ml were prepared in appropriate culture media. Cells were kept on ice until implantation. Fur was removed from the surgical site and mice were weighed prior to surgery. Mice were anaesthetised and transferred to a stereotactic frame. A midline incision was made on the head (approximately 0.5 cm) using a scalpel and a microdrill on the stereotactic frame aligned to the injection site using coordinates adjacent to the bregma to inject into the subventricular zone. A small hole was drilled in the skull using the microdrill at the injection site. 2 ml (2×10^4^ cells total) of the cell suspension was injected at a depth of 2.4 cm from the cranial surface using a 10 ml microsyringe with 28G removable needle (Hamilton, #7635-01) attached to the stereotactic frame at a flow rate of 200 nl/minute. Once injected, the needle was left in place for 4 minutes before being gradually withdrawn, and the area flushed with sterile PBS. Bone wax was applied to prevent extracranial growth of cells and the incision sutured using 5-0 reabsorbable suture. Mice received both pre-and post-surgical analgesia (Buprenorphine subcutaneous injection and oral Carprofen for the first 48 hours following surgery). Mice were monitored twice daily for the first 48 hours following surgery and once daily thereafter. Tumours were left to develop for time periods specified within results, otherwise mice were culled at the indication of neurological symptoms (ataxia, lack of reach response, balance defects, lack of grooming, loss of weight, hunched stature, or reduced movement). Animals that failed to fully recover from surgery, displayed uncontrolled pain, or which had significant surgical complications were humanely culled. The same protocol was followed for mice undergoing ‘sham’ surgery, with 2 μl of cell free media being injected. To minimise potential confounders, the same researchers performed all animal observations, delivery of any treatment, and determining humane end points.

### Generating 005-GSC FAK knockouts

FAK gene deletion of 005-GSCs was conducted using CRISPR ribonuclear proteins (RNPs) were made using IDT Alt-R^TM^ CRISPR-Cas9 system. 20 nM of trans-activating CRISPR RNA (tracrRNA) and 2 nM Alt-R CRISPR-Cas9 crRNA guide RNAs (key resource table) were annealed using a thermocycler at the following cycle settings; 95 °C for 5 minutes, ramp down to 25 °C at 5 % rate, ramp down to 4 °C at 25 % rate. Two crRNAs were selected for each round of nucleofection and separately annealed to tracrRNAs. For Cas9 control 005-GSCs, sterile water was added in place of crRNA. 10 mg of IDT Cas9 protein was added to annealed tracrRNA:crRNA complex and incubated for 10 minutes at room temperature before being stored on ice until addition to cell suspension. 4 x10^5^ 005-GSCs were resuspended in 20 ml of Lonza SE cell line 4D-Neucleofector buffer. Both tracrRNA:crRNAs were combined to form one complex containing two different crRNAs and added to cells with 10 nM of IDT electroporation enhancer. The contents were transferred into one well of a Nucleocuvette strip and electroporated using a 4D-Nucleofector X Unit (Lonza, #AAF-1003X) using a pre-optimised programme (EW-113). 80 ml of culture media was added to the well and the contents extracted and transferred to a culture flask. Cells were left to recover for approximately 1 week before lysates were generated to assess knockout efficiency.

### Spheroid assay

005 GSC cells (2,000 cells per well) were grown in U-bottom Ultra-Low Attachment plates (Costar, 7007) in standard growth media supplemented with 5 µg/mL laminin and centrifuged 200xg for 5 minutes. After 72 hours, cells were treated with 0.25 µM VS-4718 or DMSO as vehicle. 5 days after treatment, spheroids were incubated with calcein blue-AM (ThermoScientific, C1429, 2.5 µM) and propidium iodide (Sigma-Aldrich, P4170, 1 µM) for 1 hour before imaging on ImageXpress Micro XL and analysed with MetaXpress software (Molecular Devices, UK).

### Western blots

#### 005-GSC 2D western blot

1×10^5^ cells were plated into 6 well plate and cultured for 3 days. Supernatant containing non-adhered cells were collected, alongside the adhered cells using a cell scraper and combined. Cells were washed 3 times in ice cold PBS until lysis with RIPA buffer. General western blot protocol was performed as detailed previously ^83^.

#### 005-GSC spheroid western blot

005 GSC cells (3,375 cells per spheroid) were grown in MicroTissues® 3D Petri Dish® micro-mold spheroids (Sigma-Aldrich) in 24-well plates in standard growth media, following the manufacturer’s instructions. After 72 hours, spheroids were treated for 4 hours with 0.25, 0.5 and 1 µM VS-4718 or DMSO as vehicle. Westerns were performed as previously described ^83^. Antibodies used for protein visualisation are detailed in the Key Resources Table.

### FAK inhibitor treatment in vivo

VS-4718 (Selleckchem, #S7653) was prepared by dissolving the drug in sterile 0.5% hydroxypropyl methyl cellulose (HPMC) (Sigma-Aldrich, #H8384). Treatment commenced at day 10 post intracranial implantation unless otherwise stated. Mice were dosed by weight twice daily on weekdays (75 mg/kg) and once daily on weekends (150 mg/kg) via oral gavage (100 ml volume per dose) for either 7 days (tissue collection studies) or 3 weeks (survival studies). Mice in control groups received 100 ml of 0.5% HPMC via oral gavage using the same dosing schedule as treated animals. Mice were randomised at point of treatment initiation to ensure equal average body weight between the groups.

### Mouse tissue dissociation

At day 17 post intracranial implantation (unless otherwise stated) mice were culled by cervical dislocation. Whole blood was collected from heart using a heparin coated syringe. Brain, femurs and spleen tissues were harvested into chilled PBS. Brain and spleen were weighed. The non-tumour containing hemisphere, cerebellum, olfactory bulbs and spinal cord were discarded. The tumour site was excised from the right hemisphere using the remanence of needle site in the cerebrum and weighed. Tumour-containing brain tissue was mechanically disassociated using a scalpel and incubated at 37 °C in a shaking incubator (180 rpm) in DMEM containing collagenase D (2 mg/ml) (Roche, #11088858001) and DNAse (0.1 mg/ml) (Roche, #1010415001) for 30 minutes. Dissociated tissue was washed in chilled PBS + 1% BSA. Spleen tissue was mechanically disassociated using a scalpel. Bone marrow was collected from femur by removing the head of the bones and centrifuging at 2000 xg for 30 seconds. Brain, spleen and bone marrow cells were washed in chilled PBS + 1% BSA and passed through a 70 mm strainer. All tissues were subject to RBC lysis according to BioLegend (#420302) protocol. Spleen, bone marrow and blood cells were resuspended in cold PBS + 1% BSA and stored at 4 °C while performing myelin removal on brain cells. Brain cells were subject to myelin removal using magnetic labelling and LS columns/MACS Separator according to the manufacturer’s protocol (Miltenyi Biotech #130-096-433). Unlabelled, myelin-free cells were collected and all tissues were washed into cold PBS (protein free).

### Flow cytometry of tissues

Following tissue dissociation approximately 1×10^6^ cells were aliquoted in 5 ml tubes and stained with FixableBlue viability dye (eBiosceince, Thermo Fischer Scientific), before being washed into PBS + 1% BSA. Cells were incubated with Fc Blocking antibody (Biolegend). Antibodies used for immune profiling are detailed in Supplementary Table.4, and master mixes were prepared before incubation with cells. After washing, cells were fixed and permeabilised overnight using FoxP3/transcription factor staining buffer set following the manufacturer’s instructions (eBiosceince, ThermoFischer), before staining with intracellular antibody master mixes. Finally, cells were washed before being acquired on BD Biosciences LSR Fortessa^TM^ X-20 analyser, with compensation performed using AbC^TM^ Total Antibody Compensation Bead Kit (Invitrogen, #A10497) and and ArC^TM^ Amine Reactive Compensation Bead Kit (Invitrogen, #A10346). Gating and analysis of flow cytometry data was conducted using FlowJo (Version 10.10, Tree Star), with example gating strategies demonstrated in supplemental figures.

### 005-GSC Conditioned media collection and CCL2 ELISA

005-GSC cells were cultured for 3 days in 6 well plates. For cell lines treated with FAKi (VS-4718), it was added to the media during plating at a concentration of 300 nM. 0.1% DMSO was added into control flasks. On day 3 conditioned media (CM) was collected from flasks, centrifuged and filtered through a 0.22 mm sterile filter to remove any debris. Conditioned media was stored at -70 °C prior to use. CM was concentrated using Pierce protein concentrator vials (Thermo, #88515) and centrifuged at 4000 xg until the volume had reduced by half (2X concentrated). CCL2 mouse ELISA kit was purchased from Biolegend. Manufacturer’s instructions were followed, with CM samples added in triplicate. Absorbance was read at450 nm and reference wavelength of 570 nm. Concentration of secreted factors were adjusted according to concentration factor.

### Isolation of primary CD8 T cells and monocytes from mice

#### CD8 T cells

Axillary lymph nodes, inguinal lymph nodes and spleen were harvested from C57BL/6 mice. Tissues were minced using scalpel and passed through a 70 μm strainer pre-washed with T cell media (RPMI supplemented with 10% FBS, 1% Glutamine and 0.5% PS) using a syringe plunger. Flowthrough was collected and centrifuged at 300 xg for 5 minutes. Cells were washed again into ice cold PBS, before red blood cell lysis following the manufacturer’s instructions (Biolegend, 420302). Splenocytes were then washed into PBS+ 1% BSA before counting, and CD8+ T cell isolation using CD8a+ isolation kit (Miltenyi, 130-104-075) following the manufacturer’s instructions. Purified CD8+ T cells were then used for further assays detailed below.

#### Monocytes

Fore and hind limbs were harvested from female C57BL/6 mice under sterile conditions. Femurs, tibias and humeri were stripped of tissue and the ends snipped using a scalpel, placed into 0.5 ml tube inside a 1.5ml tube at centrifuged at 2000 xg for 1 minute to flush out the bone marrow, and washed in PBS+ 1% BSA. Harvested bone marrow was passed through a 40 mm cell strainer using PBS+ 1% BSA and centrifuged at 300 xg for 5 minutes. Bone marrow was washed into PBS containing 0.5% BSA and 2 mM EDTA, for monocyte isolation using Miltenyi Biotec monocyte isolation kit (BM), mouse (#130-100-629) following the manufacturer’s protocol. Isolated monocytes were cultured in DMEM – high glucose (Sigma-Aldrich, #D6429) supplemented with 10% heat inactivated FBS, 0.5% pen-strep, 5 ng/ml recombinant mouse GM-CSF (R&D, #415-ML) and 2.5 ng/ml recombinant mouse IL-4 (Biolegend, #574302) for a maximum of 5 days.

### CD8 T cell in vitro stimulation assay

CD8^+^ T cells were isolated from C57BL/6 mice as detailed above. 50,000 cells were plated into a 96-well plate and stimulated with plate bound anti-CD3 (7.5μg/ml) (Biolegend, 100340), with addition of IL-2 (0.005μg/ml) (Biolegend, 575406) and anti-CD28 (2μg/ml) (Biolegend, 112116), either in 100% T cell media, or in the presence of 30% CM from 005-GSC cells as detailed above. T cells were stimulated for 3 days and then harvest from the plate for analysis by flow cytometry as detailed above. For proliferation analysis, at point of CD8+ isolation, cells were incubated with 5nM CellTrace Violet as per the manufacturer’s instructions (Thermo Fischer Scientific, C34557), and analysed by flow cytometry.

### CD8^+^ T cell and bone marrow derived macrophages co-culture assay

#### CD8+ T cells

CD8^+^ T cells were isolated as detailed above, and stimulated overnight in a flask pre-coated with anti-CD3 (7.5μg/ml) (Biolegend, 100340), with addition of IL-2 (0.005μg/ml) (Biolegend, 575406) and anti-CD28 (2μg/ml) (Biolegend, 112116) to media.

#### Bone marrow derived macrophages (BMDMs)

Hindlimbs were isolated from C57BL/6 mice and the muscle stripped. The femur and tibia were separated and the ends of each bone were cut to create open ends. The bone marrow was subsequently flushed out of the bones with DMEM-Hi media (Life technologies, D6429) using a 25G needle into a 70μm cell strainer (Falcon, 352350) and cells were washed in DMEM-Hi at 300 xg for 5 minutes. The bone marrow cells were resuspended into a volume of 1200μl of which 200μl were added to a well of a 6 well plate containing 2 ml of macrophage differentiation media (DMEM-Hi + 20 ng/ml m-CSF (Biolegend, 576406)). The cells were then left in culture for 4 days to differentiate into macrophages.

#### Co-culture

Macrophage media was aspirated, and BMDMs were washed twice with warm PBS, before detachment with TrypLE and cell scraping. After washing macrophages were resuspended in co-culture media (T cell media + 20 ng/ml m-CSF) and counted. T cells were resuspended in the flask before being collected and centrifuged. They were then washed with PBS, resuspended in co-culture media and counted. Macrophages and T cells were then plated into a 96 well plate in the ratios described in figure legends, and left for 6 days before analysis by flow cytometry.

### Ms4a3^Cre^-Rosa^TdT^ and Cdh5-CreER^T2^; Rosa26^lsl-tdT^ multiphoton brain slice image collection

#### Tissue processing

Brains were collected into 4% paraformaldehyde for fixing. After 24 hours, tissue was stored in PBS, embedded in plastic molds with 2% low-melting point agarose and sectioned close to the injection region with 300 mM thickness using a Vibratome Leica VT1200 S.

#### Multiphoton imaging

Two-photon fluorescence images were acquired using a custom-built multi-modal microscope setup. A picoEmerald S (APE, Berlin, Germany) laser provided both a tunable pump laser (set to 797.4 nm, 2 ps, 80 MHz repetition rate) and a spatially and temporally overlapped Stokes laser (1031 nm, 2 ps, 80 MHz repetition rate). The output beams were inserted into the scanning unit of an Olympus FV1000MPE microscope using a series of dielectric mirrors and a 3× lens-based beam expanding module. The resulting 3.6 mm beams were expanded by a further 3.6× within the microscope and directed into an Olympus XLPL25XWMP N.A. 1.05 objective lens using a short-pass 690 nm dichroic mirror (Olympus). Backscattered two-photon fluorescence were separated from backscattered excitation light using a short-pass 690 nm dichroic mirror and IR cut filter (Olympus). A series of filters and dichroic mirrors were then used to deconvolve the different emission signals onto two photomultiplier tubes (PMT). Emissions were filtered using: FF552-Di02, FF483/639-Di01 and FF510/84 for GFP and tdTomato signals using FF552-Di02, and FF440/520-Di01 and HQ610/75m (Chroma). All filters are from Semrock unless marked otherwise. Laser powers after the objective were measured up to 70 mw for the pump and Stokes laser. Images were recorded by FV10-ASW software (Olympus). Images were aquired using 512 × 512 or 1024 × 1024 pixels over a 509 × 509 μm field of view, with a pixel dwell time of 4 μs. Whole brain slices were imaged using the multi-area time-lapse function, whilst image stacks of 155 slices were recorded with a z interval of 1 μm, PMT voltages were adjusted with depth to maintain brightness.

#### Image processing

Image processing was done in FIJI (1.54r) ^84^. Stitched images were thresholded to produce a mask of the total brain area, whilst tumour masks were drawn manually from the same image. A ‘brain only’ area was produced by subtracting the tumour mask from the brain mask. TdTomato area was calculated by first using the Image Calculator function to subtract the GFP image, removing any fluorescence bleed-through. The TdTomato area was then measured by thresholding and using the tumour and ‘brain only’ masks as area masks.

### Measurement of tumour vasculature

Cdh5-CreER^T2^; Rosa26^lsl-tdT^ mice ^41,42^ were injected intracranially with 005-GSC cells and treated with vehicle (HPMC) or VS-4718 as described above. After 7 days treatment mice were culled and tumour slices imaged to visualise the tdTomato labelled endothelial cells as detailed above. Images were segmented using the Labkit addon ^85^ in FIJI ^84^. The classifier (default settings) binarized the entire image into background and vessels. Opening and closing was then applied to the images to remove noise and fill in small holes and centrelines were extracted as a graph using Kimimaro (Supplementary Figure 3G). From the centrelines total vessel length per tissue area was determined as a measure of vascular density (Supplementary Figure 3H).

### Monocyte VCAM-1 adhesion assay

96-well plates were coated with 5 μg/ml VCAM-1 (rmVCAM-1/Fc Chimera R&D, #643-VM) diluted in PBS at 4 °C overnight. Monocytes were isolated from female C57BL/6 mice as described above. VCAM-1 was aspirated and non-specific binding was blocked with DMEM 10% FBS. Monocytes were adjusted to 20,000 cells per 100 ul in suspension. 300 nM VS-4718 was added to cell suspension and incubated at 37 oC for 30 minutes. For anti-α4β1 blocking antibody 10 μg/ml of both clone R1-2 and clone 9C10 (Biolegend) were added to cell suspension and incubated for 37 °C for 30 minutes. After incubation, monocytes were seeded onto the pre-coated plates and left to adhere for 30 minutes at 37 °C. Media and non-adhered cells were removed, wells were gently washed with PBS and remaining adherent cells were fixed with 4 % PFA for 15 minutes at room temperature. 4% PFA was removed and wells washed with PBS. 0.05 % crystal violet solution was added to wells and incubated for 30 minutes at room temperature. Wells were washed twice with H_2_O, left to dry, then representative images were captured using Incucyte SX5, with 5 images taken per well.

### Adoptive transfer of sgFAK Ms4a3^Cre^-Rosa^TdT^ monocytes

#### Tumour establishment

Male C57BL/6 mice were injected with 005-GSC cells intracranially as described above. Tumours were left for 15 days.

#### FAK knockout of monocytes

Monocytes were harvested from male Ms4a3^Cre^-Rosa^TdT^ mice as described above and pooled. CRISPR-Cas9 directed deletion of FAK using guide RNAs (sgFAK) was conducted as described above using the IDT Alt-R^TM^ CRISPR-Cas9 system. For nucleofection of RNPs, P3 cell line 4D-Neucleofector buffer (Lonza) was used on 4D-Nucleofector X Unit using a pre-optimised programme (CM-137). Monocytes were then resuspended in warm monocyte media and left in 37 °C 5 % CO_2_ incubator to recover overnight before adoptive transfer. Adoptive transfer: At day 15 post tumour injection, sgFAK and sgCTRL monocytes were collected and washed into PBS with no Mg^2+^ and no Ca^2+^ to prevent clumping. Monocytes were resuspended at a concentration of 1×10^6^ cells per 100 μl and 100 μl was delivered intravenously into 005-GSC tumour bearing mice. FAK expression was checked by flow cytometry as detailed above, using anti-FAK antibody (Thermo Fischer Scientific, #MA5-41093) conjugated to CoraLite® Plus 647 with FlexAble CoraLite® Plus 647 Antibody Labeling Kit for Rabbit IgG (Proteintech, #FA003-50), following manufacturer’s instructions. 005-GSC FAK-NULL cells were used as a negative anti-FAK antibody control to set positive FAK staining gate.

#### Tissue harvest and flow cytometry

After 4 days post adoptive transfer, mice were culled and brain, spleen, bone marrow and whole blood were collected as described above. For controls a 005-GSC tumour bearing mouse with no adoptive transfer, and a mouse with no tumour burden but received adoptive transfer was also collected. Tissues were processed for flow cytometry as described above, however half of the cells per sample were not fixed to allow for analysis of TdTomato signal. Flow cytometry analysis was conducted as detailed above.

### FAK and pFAK immunohistochemical staining of brain tumour patient tissue microarray (TMA)

A TMA of brain tumour patients was used which has been detailed previously ^65^. Formaldehyde-fixed tumour samples were embedded in paraffin, sectioned onto slides. Samples were rehydrated with xylene twice and dewaxed with decreasing gradients of ethanol. For antigen retrieval, samples were treated with 0.1 M citrate buffer (pH 6.0) at 100°C and blocked with peroxidase blocking solution (Agilent, Santa Clara, CA, USA) for 5 minutes and then with serum-free protein solution (Agilent) for 10 minutes at room temperature. Samples were incubated with primary antibodies pFAK Y576 (Abcam, ab226847, 1:300) or mouse total FAK pY397 (BD Biosciences, 610087, 1:100) overnight at 4°C. Sections were washed 2 times with 0.05% v/v TBS-T, incubated with EnVision System HRP labelled polymer anti-rabbit (Agilent) for 30 minutes, washed twice with 0.05% v/v TBS-T and exposed to DAB+ (Agilent) for 5 minutes. Next, samples were counterstained with Mayer’s hematoxylin (Sigma-Aldrich) for 2 minutes and exposed to increasing gradients of ethanol and xylene before mounting with DPX (Sigma-Aldrich). Immunostained tissue slides were digitised using NanoZoomer slide scanner (Hamamatsu, Shizuoka, Japan) with 40X magnification.

### Scoring of FAK and pFAK of brain tumour TMA

#### pFAK

Of 195 cores across the pFAK-stained TMA, 108 were evaluable for pFAK expression (N=87 not evaluable: N=64 missing, N=10 necrosis, N=6 no identifiable tumour, N=6 affected by substantial artefact, N=1 poor imaging quality). 81 of the evaluable cores (75.0%) were negative, 12 (11.1%) demonstrated weak positivity, and 15 (13.9%) demonstrated moderate or strong positivity. The 108 evaluable cores represented samples from 42 unique brain tumour patients, of which 27 had a glioblastoma (GBM) diagnosis and were included in subsequent analyses. All cases had samples from primary diagnosis; 8 also had evaluable samples from material taken at recurrence. Per-patient pFAK status was determined using the samples from diagnosis: individuals were considered pFAK positive if any evaluable TMA cores contained at least weak tumour cell pFAK positivity. If all evaluable cores from diagnosis were negative for pFAK expression, cases were considered to be pFAK negative. Of the 27 evaluable GBM cases, 11 (40.7%) were pFAK-positive and 16 (59.3%) were pFAK negative.

#### FAK

Of 195 cores across the FAK-stained TMA, 105 were evaluable for FAK expression (N=90 not evaluable: N=74 missing, N=8 necrosis, N=4 no identifiable tumour, N=4 affected by substantial artefact). The evaluable cores represented 43 unique brain tumour patients, of which 28 had a glioblastoma diagnosis. 27 of these represented cases with evaluable material from diagnosis and were included in subsequent analysis (n=1 with recurrence samples only); 7 cases had matched evaluable samples from diagnosis and recurrence. Per-patient FAK expression was quantified using the samples from diagnosis: expression of FAK in evaluable cores was quantified using histoscore, whereby the percentage of tumour cells (0-100%) were multiplied by their expression intensity (0, negative; 1, weak positive; 2 intermediate positive; 3, strong positive) to produce a quantified score from 0-300 ^86^. Both nuclear-specific and cytoplasmic-specific histoscores were calculated. Where multiple cores were evaluable for FAK expression per case, a patient-level histoscore was calculated as the mean histoscore across cores.

### Stratification of TMA with FAK histoscores and matched RNAseq analysis

RNA was extracted from 4 µm FFPE sections from the cores used to generate the TMA as detailed above. RNA-sequencing analysis was carried out by Genomics Core Technology Unit (Genomics CTU, Belfast, Queen’s University Belfast). RNA-sequencing data (for 29 patients) were matched against evaluable FAK histoscores and were normalised using DESeq2^87^. For inference of bulk transcriptional state, gene set variation analysis (GSVA^88^ was performed using the mesenchymal, proneural and classical subtype reference gene sets as described by Wang *et al.*^66^. Cell type inference was performed using the GBMDeconvoluteR^67^ and resulting arbitrary cell type abundance scores were scaled using Z-scoring to yield a Scaled Abundance Score. Data were plotted using ComplexHeatmap^89,90^ using R version 4.5.1.

### Correlation of FAK and pFAK expression with CD163 and CD68 expression on TMA

pFAK expression status and FAK histoscores were compared against quantified levels of CD163- and CD68-positive cells available from a previous study ^65^. Correlation of continuous variables was assessed using Spearman’s rank correlation coefficient (Spearman’s rho). Differences in CD163 and CD68 levels between categorical groups (pFAK positive vs pFAK negative) was assessed using Mann-Whitney U tests.

### Statistical analysis

Statistical analyses were performed using either Excel (Microsoft Office 2021), GraphPad Prism version 10.5 (Dotmatics), R version 4.4.0 or above. To determine differences between groups, unpaired t-tests were performed for normally distributed data, otherwise Mann-Whitney U test was used, unless otherwise stated in figure legends. Multiple comparisons (with corrections) were conducted with one-way ANOVA or Kruskal-Wallis test, unless otherwise stated in figure legends. Kaplan-Meier methods were used for plotting of survival curves with log-rank tests used to assess significant differences between groups. Each mouse was considered an experimental unit. Individual mice were excluded from survival analysis if were culled within the experimental timeframe due to non-tumour related symptoms, and/or no tumour present on histological analysis. Individual mice were excluded from flow cytometry analysis if the intracellular staining had been determined to have been unsuccessful or contamination of samples with large amounts of debris (myelin). P values are displayed as: *= <0.05, ** = <0.01, *** = <0.001 and **** = <0.0001.

## Author contributions

Conceptualization, E.R.W, M.C.F and V.G.B.;

Methodology, E.R.W, A.B, G.C, M.F, R.L.H, A.E.P.L, R.E, T.W, J.A.A, B.P, M.M, M.L.;

Formal analysis, E.R.W, A.B, G.C, R.L.H, A.E.P.L, J.C.O, R.E, S.A.S, T.W, J.A.A, T.B, B.P, M.L.;

Investigation, E.R.W, A.B, G.C, R.L.H, M.F, A.E.P.L, J.C.O, S.A.S, T.B, M.H, B.P, M.M, F.L, M.L.;

Resources, R.E, S.A.S, C.S, T.R.H, P.M.B, Z.L, F.G, M.O.B, A.S, R.G.;

Writing – original draft, E.R.W. V.G.B;

Writing – review & editing, all authors;

Visualization, E.R.W, A.B, G.C, R.H, M.F, A.E.P.L, M.L.;

Supervision, E.R.W, A.R, J.A.A, T.R.H, P.M.B, M.O.B, R.G, M.C.F and V.G.B.;

Project administration and funding acquisition, M.C.F and V.G.B.

## Acknowledgements

We would like to thank the following service providers at the Institute of Genetics & Cancer, University of Edinburgh; Histology Research Service for tissue microarray preparation and mouse tissue sample preparation for IHC, and the Flow Cytometry Facility for running samples. We thank the Lothian NRS Bioresource for access to the tissue microarray (REC approval 15/ES/0094). We also thank Mahir Haque for assistance with performing *in vitro* flow cytometry analysis of monocytes, and Dr Vanessza Fentor for help organising the patient RNAseq data. We would like to thank Prof Samuel Rabkin (Harvard University, USA) for providing the 005-GSC cells, and Prof. Peter Angel, (German Cancer Research Centre, Heidelberg, Germany) for providing the mGB2 cells. We thank Prof Neil Carragher (University of Edinburgh) for help drafting the manuscript.

