## Supplementary Figures 1-10 for "FAK inhibitor treatment systemically regulates the glioblastoma immune environment via blocking monocyte trafficking"

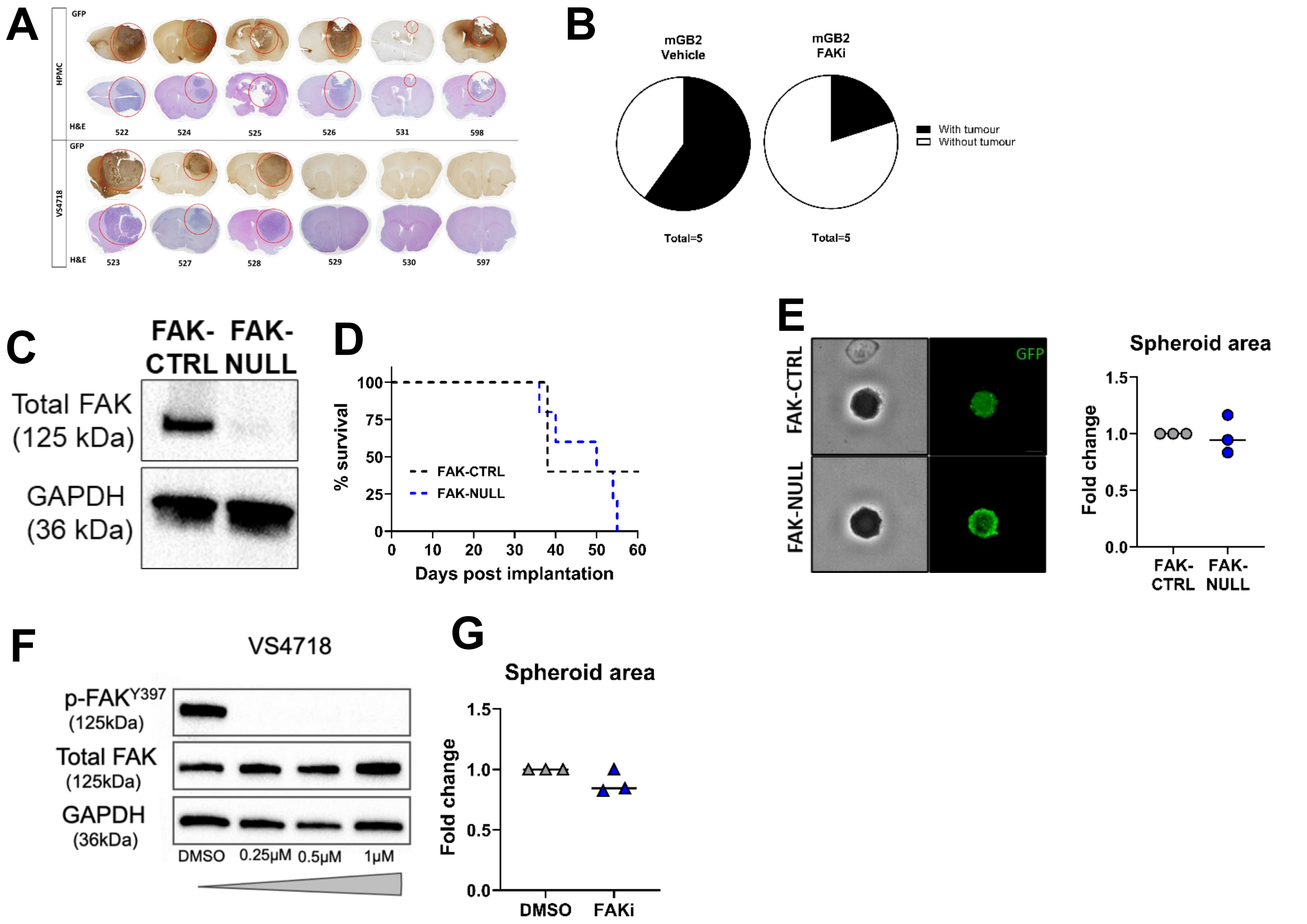


**Supplementary Figure.1**

**A)** H&E and GFP immunohistochemistry of brains taken from second experiment (survival shown in Figure.1A).

**B**) Percent of mice classed as bearing mGB2 tumours (with tumour) or without (without tumour) at 200 day end point post tumour inoculation.

**C)** Western blot showing loss of FAK protein after CRISPR-Cas9 gene deletion of FAK

**D)** Survival analysis of a second CRISPR-Cas9 FAK knockout cell line

**E)** Spheroids of 005-GSC FAK-CTRL and FAK-NULL cells were assessed for their growth (Area – GFP expression) in vitro. n=3 biological replicates, statistical significance assessed by unpaired t-tests.

**F)** Western blot of phospho-FAK, total FAK and loading control GAPDH protein in 005-GSC cells in vitro after FAKi treatment.

**G)** Analysis of 005-GSC spheroid area (GFP signal) in vitro after FAKi treatment.

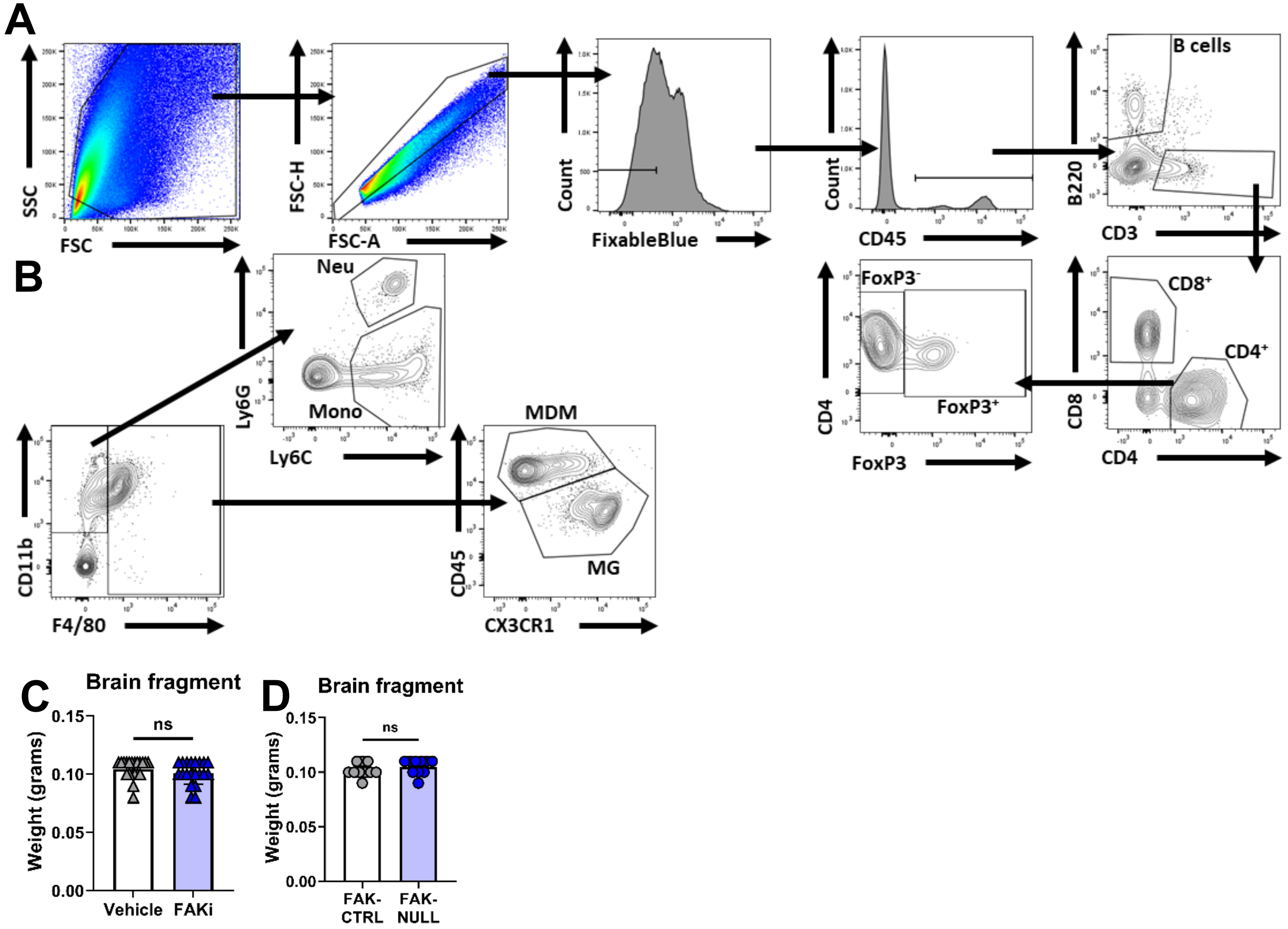


**Supplementary Figure.2**

**A)** Flow cytometry gating strategy for defining B cells, CD8$^{+}$ T cells, CD4$^{+}$ FoxP3$^{-}$ T cells and CD4+ FoxP3$^{+}$ T regulatory cells

**B)** Flow cytometry gating strategy for defining monocyte derived macrophages (MDM), microglia (MG), neutrophils (Neu) and monocytes (Mono).

**C+D)** Weight of brain fragments containing the tumours used for flow cytometry analysis.

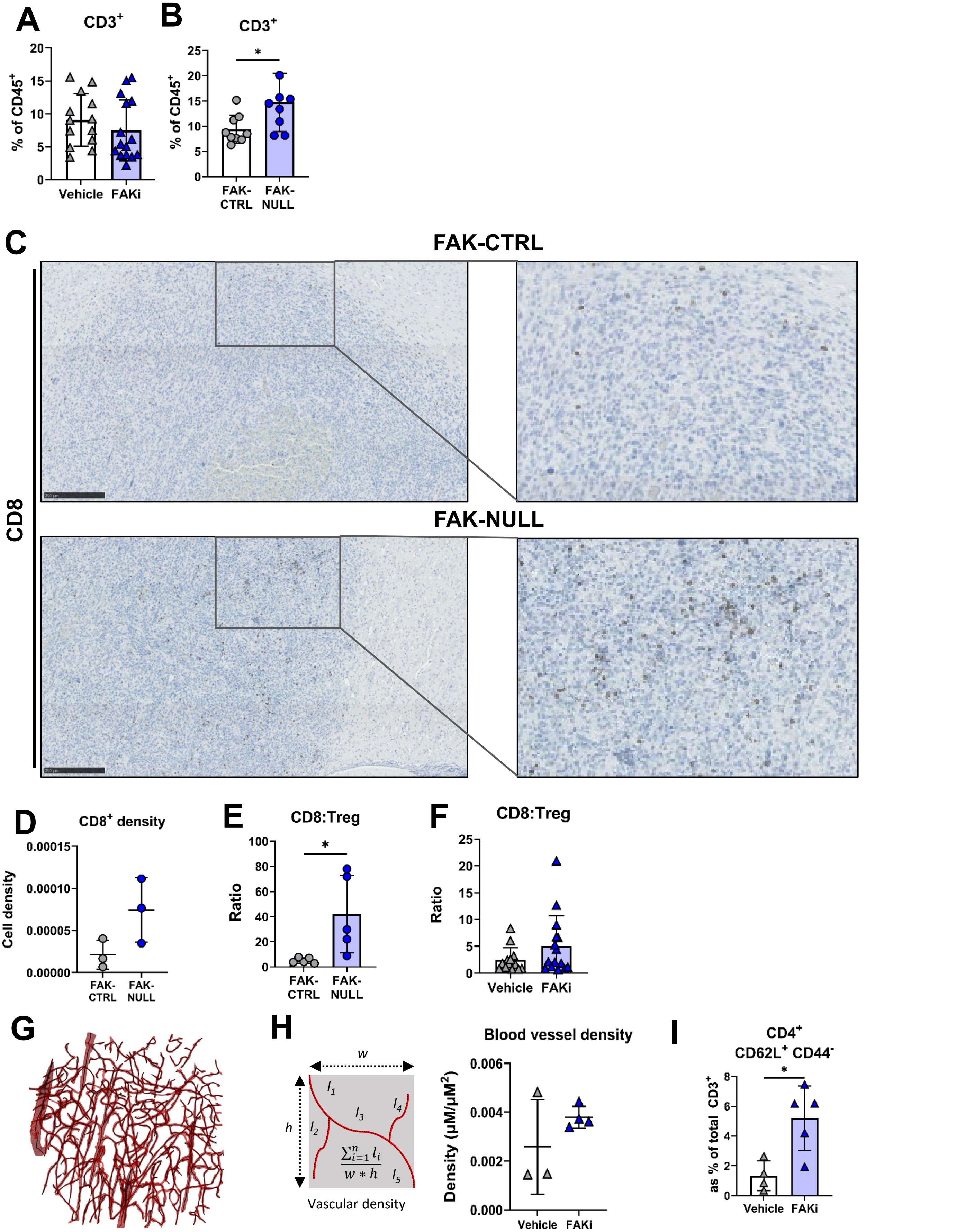


**Supplementary Figure.3**

**A+B)** Total T cell ($CD3^+$) infiltration shown as percentage of $CD45^+$ cells (as percent of total cells).

**C+D)** Example staining of $CD8^+$ T cells in 005 tumours, with quantification shown in D.

**E+F)** Ratio of $CD8^+$ T cells to Treg cells in tumours.

**G)** Representative reconstructed surface of the tumour vascular network with the centrelines in black inside the surface which were used to visualise blood vessels from which vessel density metrics were obtained.

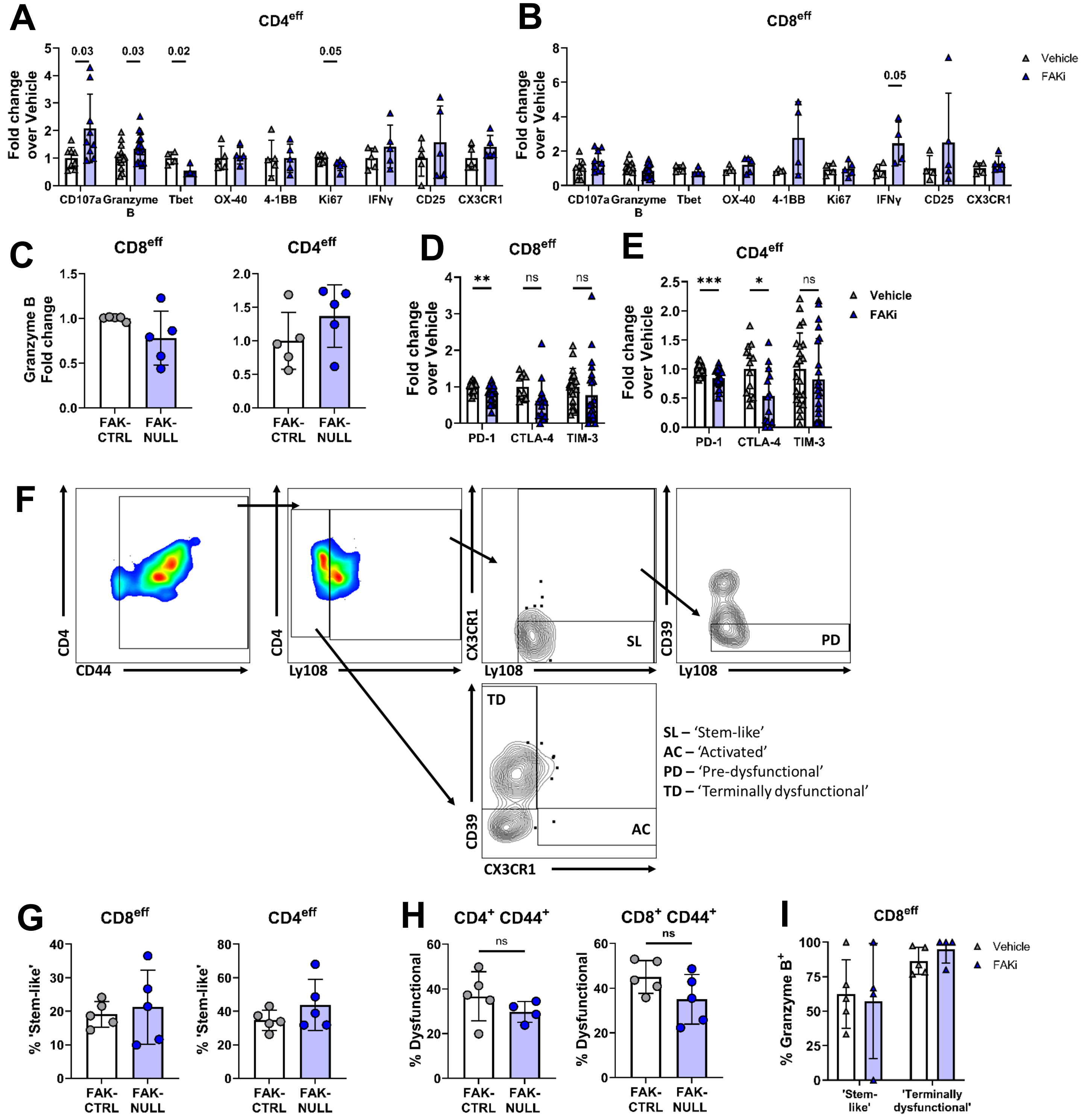


**Supplementary Figure.4**

**A+B**) Analysis of expression of key activation markers on effector CD4$^{+}$(A) and CD8$^{+}$(B) tumour infiltrating T cells. Individual mouse data shown here which was used to generate the heat map shown in Figure.2A.

**C**)Granzyme B expression in CD8eff and CD4eff in 005-GSC FAK-CTRL or FAK-NULL tumours, as fold change over mean of FAK-CTRL.

**D+E**) As in C for inhibitory receptor expression on CD8$^{eff}$ and CD4$^{eff}$ respectively.

**F**) Example gating of T cell exhaustion phenotypes.

**G+H**) Percent of CD8eff and CD4effs classified as 'stem-like', 'activated', 'pre-dysfunctional' and 'dysfunctional' in 005-GSC FAK-CTRL or FAK-NULL tumours.

**I**) Percent of 'stem-like' and 'terminally dysfunctional' CD8eff T cells expressing Granzyme B in 005-GSC tumours treated with FAKi.

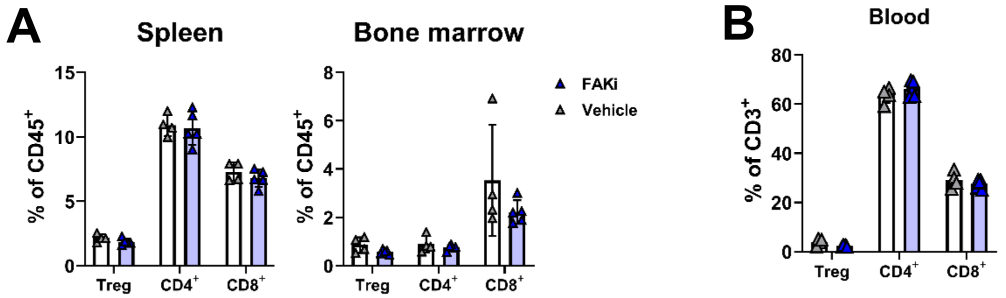


**Supplementary Figure.5**

**A**) Analysis of Treg, $CD4^+$ and $CD8^+$ populations as a percent of total $CD45^+$ cells in Bone marrow and spleen of 005-GSC tumour bearing mice treated with FAKi.

**B**) Analysis of Treg, $CD4^+$ and $CD8^+$ populations as a percent of total $CD3^+$ cells in blood of 005-GSC tumour bearing mice treated with FAKi.

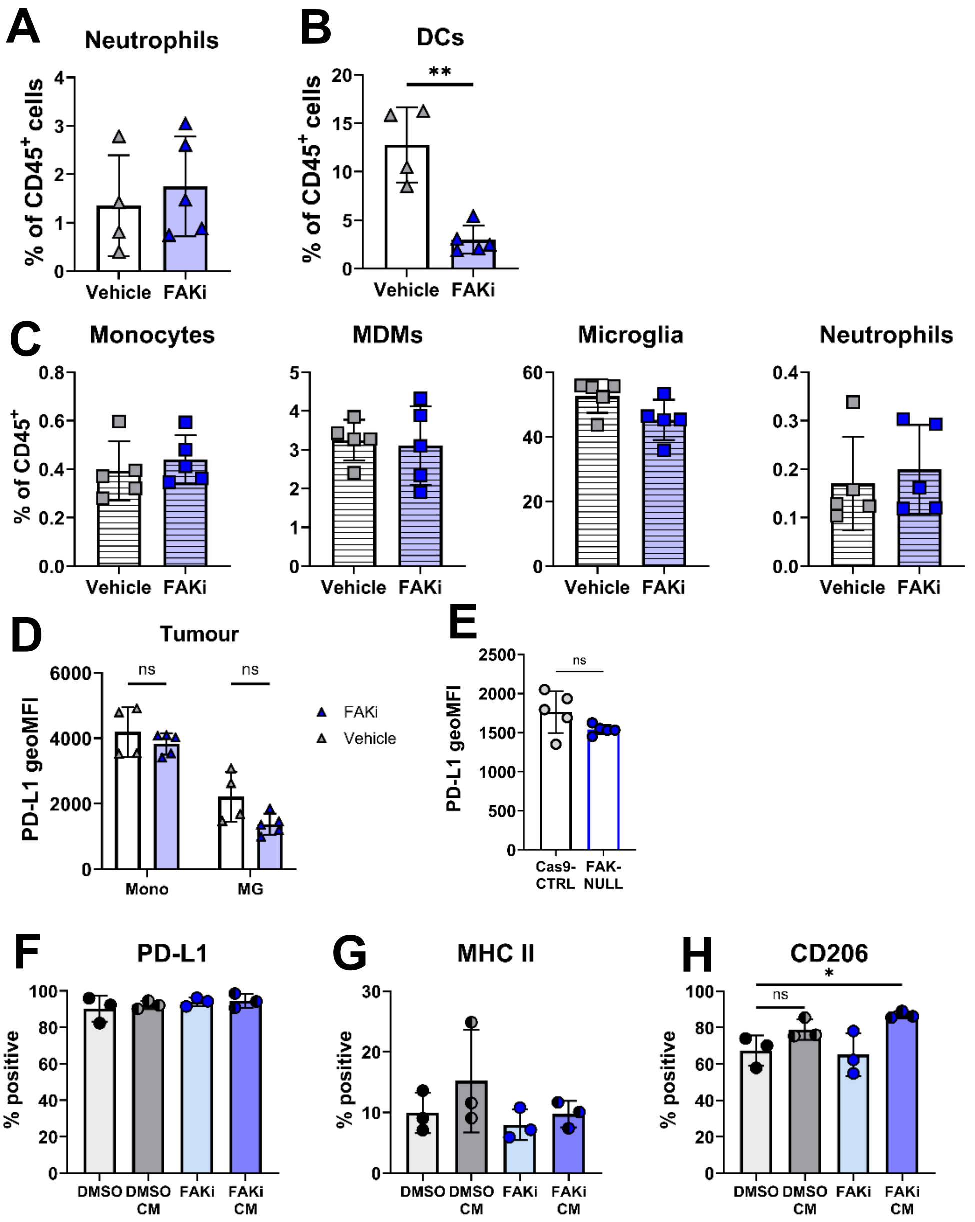


**Supplementary Figure.6**

**A**) Percent of neutrophils as percentage of total immune cells ($CD45^+$) in tumour

**B**) Percent of dendritic cells (DCs) as percentage of total immune cells ($CD45^+$) in tumour

**C**) Brain infiltrating myeloid populations as a percent of total $CD45^+$ cells in shame surgery mice treated with FAKi

**D**) Expression level of PD-L1 on tumour infiltrating monocytes and microglia in FAKi treated 005-GSC tumours

**E**) Expression level of PD-L1 on MDMs in FAK-NULL tumours

**F-H**) Percent of BMDMs which express PD-L1 (**F**), MHC II (**G**) and CD206 (**H**) in the presence of either direct FAKi addition (DMSO and FAKi) or conditioned media from FAKi treated 005-GSC cells (DMSO CM and FAKi CM).

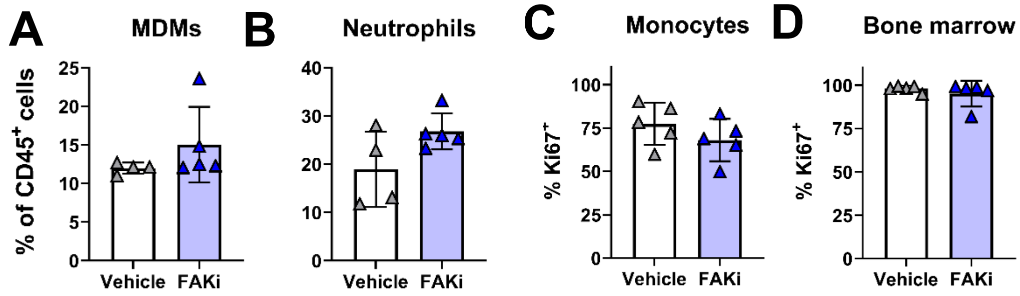


**Supplementary Figure.7**

**A+B**) Bone marrow myeloid populations as a percent of total $CD45^{+}$ cells in FAKi treated 005-GSC tumour bearing mice.

**C**) Percentage of monocytes expressing Ki67 in the tumour of FAKi treated 005-GSC tumour bearing mice.

**D**) Percentage of monocytes expressing Ki67 in the bone marrow of FAKi treated 005-GSC tumour bearing mice.

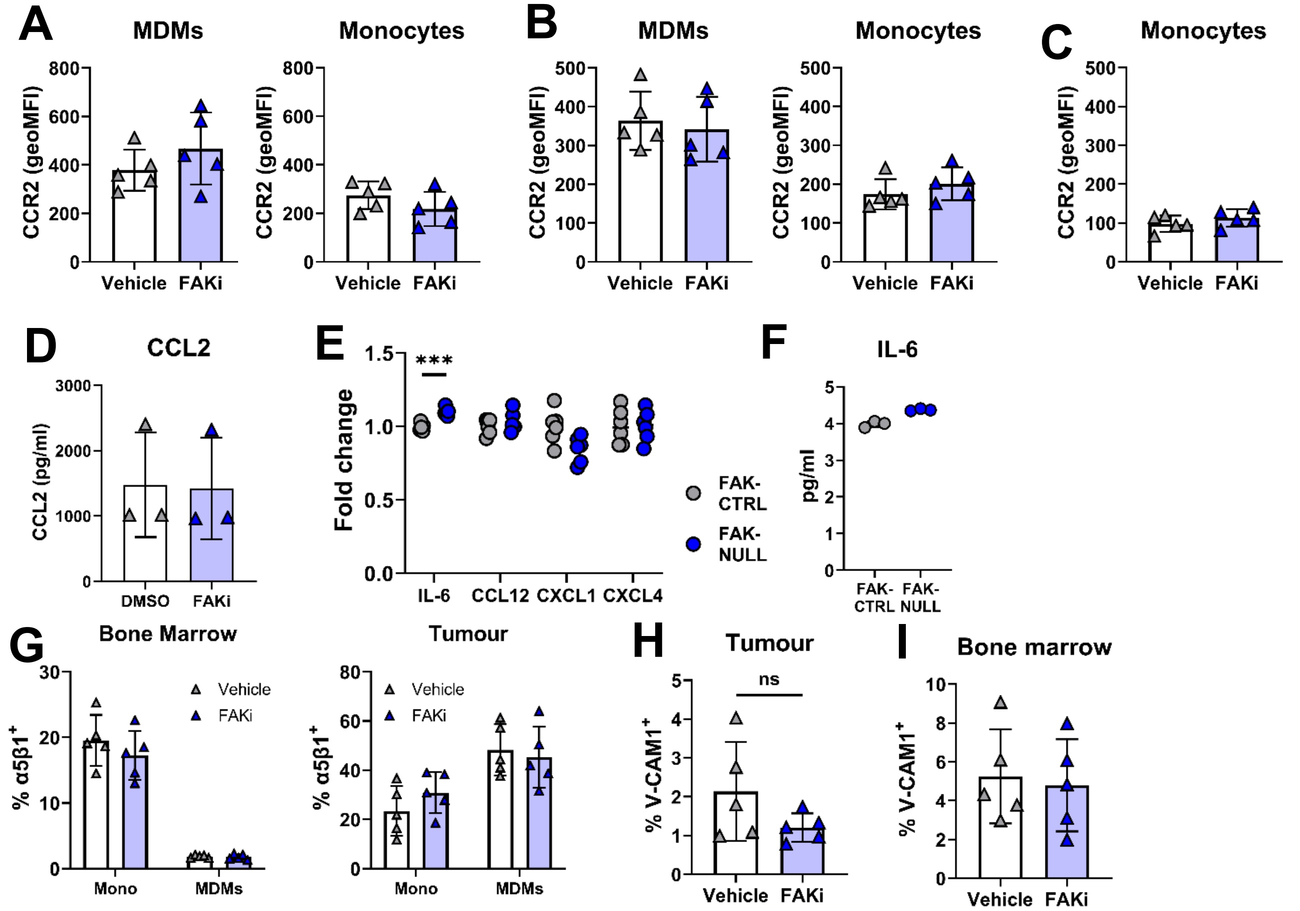


**Supplementary Figure.8**

**A**) Expression levels of CCR2 on bone marrow myeloid cells of FAKi treated 005-GSC tumour bearing mice.

**B**) As for **A** for spleen myeloid cells.

**C**) As for **A** for whole blood monocytes

**D**) Quantification of CCL2 secretion by ELISA of FAKi treated 005-GSC cells in vitro

**E**) Secretion of relevant cytokines involved in myeloid cell trafficking by either 005-GSC FAK-CTRL or FAK-NULL cells in vitro, demonstrated as fold change over control.

**F**) Absolute concentration of IL-6 being secreted in I.

**G**) Percent of monocytes and MDMs expressing α5β1 in brain and bone marrow of 005-GSC tumour bearing mice treated with FAKi.

**H+I**) Percent of endothelial cells ($CD45^-$ $CD31^+$) VCAM-1$^+$ in tumour (**I**) and bone marrow (**J**).

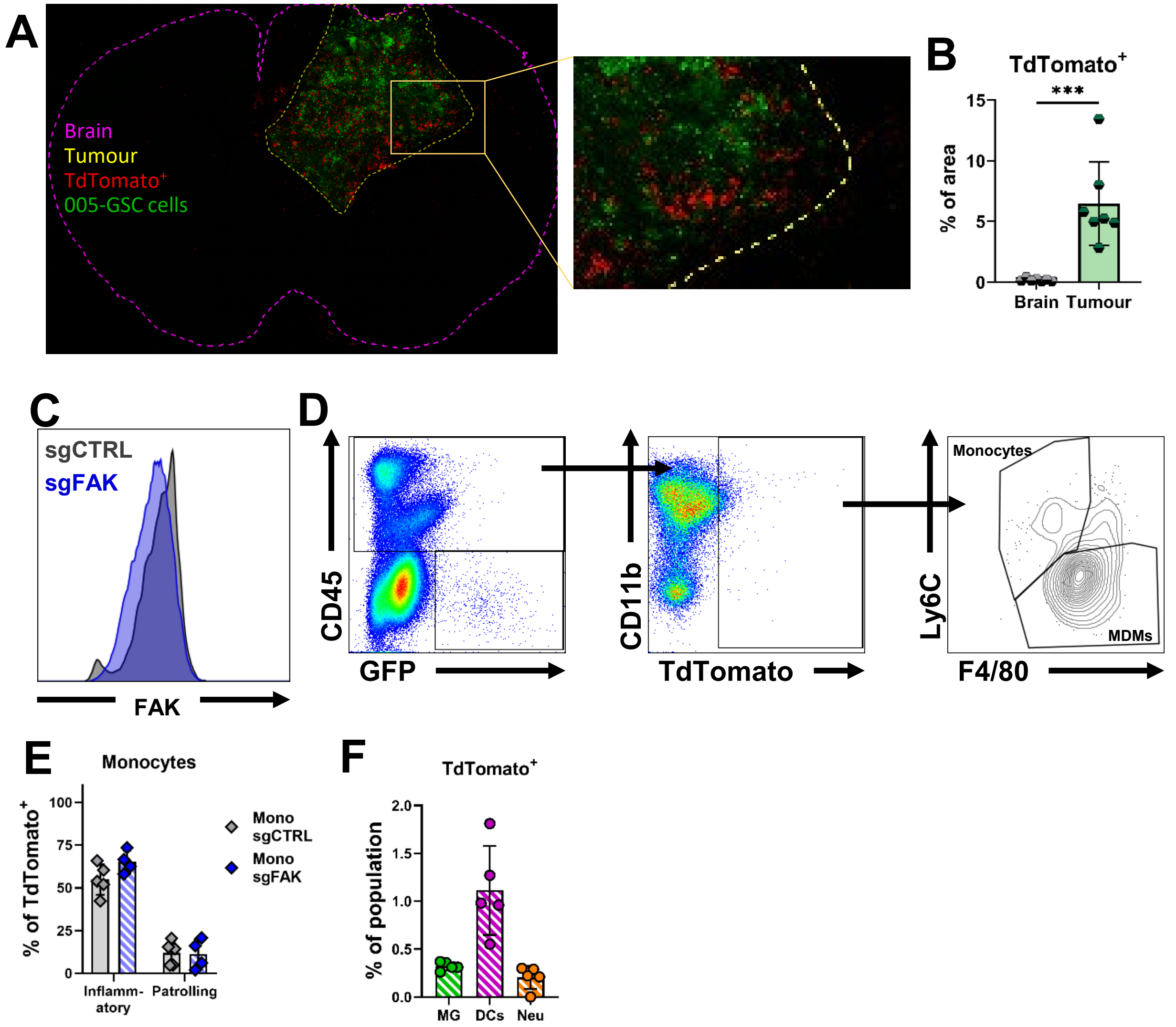


**Supplementary Figure.9**

**A**) Example of multiphoton imaging of brain slices from $Ms4a3^{Cre}$-$Rosa^{TdT}$ mice bearing 005-GSC tumours at day 20 post implantation. Brain outline is shown in purple, tumour outline shown in yellow, tumour cells are GFP$^+$ (green) and monocytes/MDMs are TdTomato$^+$ (red). Cut out (yellow box) is not to scale.

**B**) Quantification of TdTomato$^+$ cells inside or outside of tumour in A

**C**) Example of reduction in FAK staining on flow cytometry analysis of monocytes in vitro with quantification in Figure.5B

**D**) Gating strategy for flow cytometry analysis of TdTomato$^+$ cells and gating of TdTomato$^+$ monocytes and macrophages in tumours. Gates set using combination of FMOs and tissues from mice with didn't receive either 005-GSC cells (no GFP) or no adoptive transfer (no TdTomato).

**E**) Percent of TdTomato$^+$ monocytes which are inflammatory or patrolling in tumours.

**F**) Percent of non-monocyte/macrophage myeloid cells which are TdTomato$^+$ in tumours.

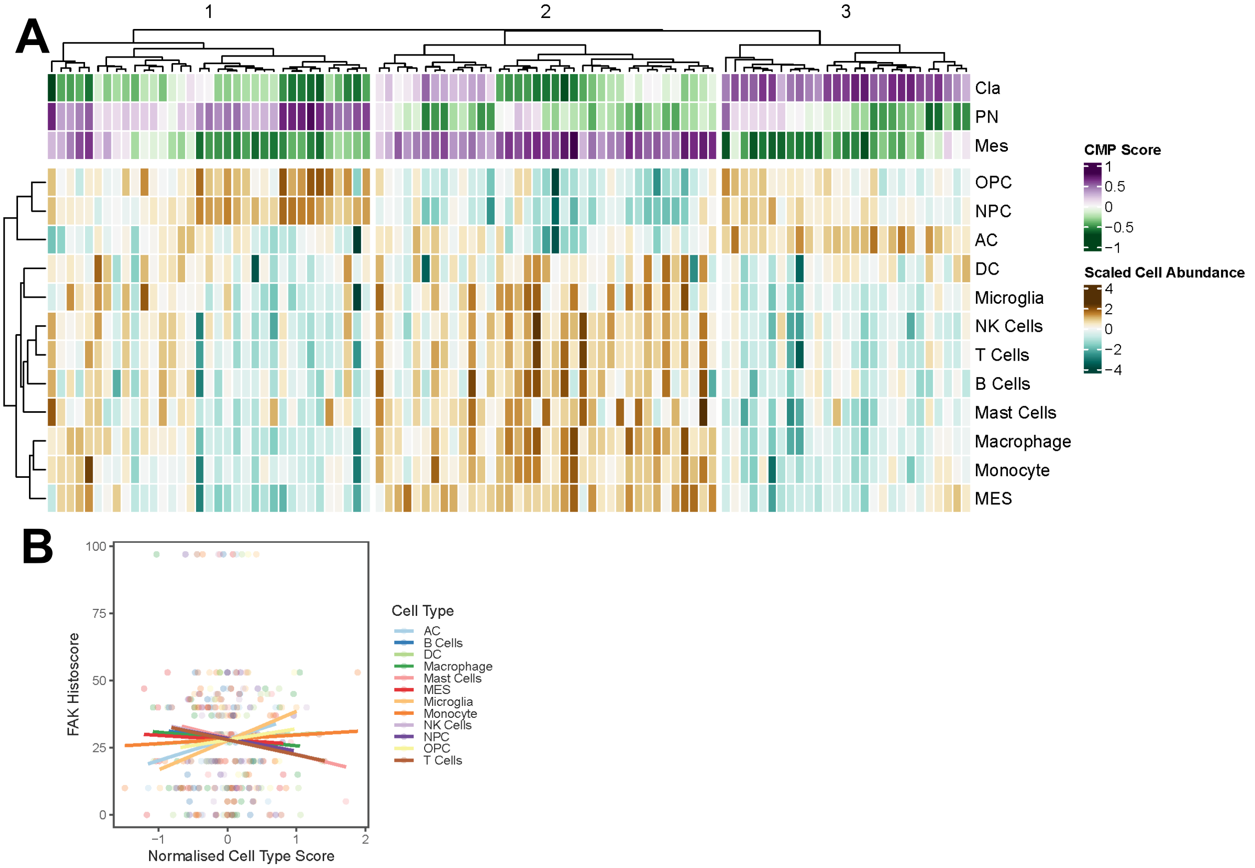


**Supplementary Figure.10**

**A)** Heatmap showing classical (Cla), proneural (PN) and mesenchymal (Mes) bulk RNA scores for 99 patient tumours collected and sequenced by the Clinical Proteomic Tumour Analysis Consortium, and inferred deconvoluted cell type abundances.

**B)** Correlation of FAK histoscores with inferred deconvolved cell type abundances for GBM TMA (shown in Figure 5I).

Immune cells: B cells, DC – Dendritic cells, Macrophage, Mast cells, Microglia, Monocyte, NK cells, T cells. Neoplastic cells: AC – astrocyte-like, MES – mesenchymal cells, NPC – neuronal progenitor-like, OPC – oligodendrocyte progenitor-like
